# Rail-RNA: Scalable analysis of RNA-seq splicing and coverage

**DOI:** 10.1101/019067

**Authors:** Abhinav Nellore, Leonardo Collado-Torres, Andrew E. Jaffe, José Alquicira-Hernández, Jacob Pritt, James Morton, Jeffrey T. Leek, Ben Langmead

## Abstract

RNA sequencing (RNA-seq) experiments now span hundreds to thousands of samples. Current spliced alignment software is designed to analyze each sample separately. Consequently, no information is gained from analyzing multiple samples together, and it is difficult to reproduce the exact analysis without access to original computing resources. We describe Rail-RNA, a cloud-enabled spliced aligner that analyzes many samples at once. Rail-RNA eliminates redundant work across samples, making it more efficient as samples are added. For many samples, Rail-RNA is more accurate than annotation-assisted aligners. We use Rail-RNA to align 667 RNA-seq samples from the GEUVADIS project on Amazon Web Services in under 16 hours for US$0.91 per sample. Rail-RNA produces alignments and base-resolution bigWig coverage files, ready for use with downstream packages for reproducible statistical analysis. We identify expressed regions in the GEUVADIS samples and show that both annotated and unannotated (novel) expressed regions exhibit consistent patterns of variation across populations and with respect to known confounders. Rail-RNA is open-source software available at http://rail.bio.

## 1 Introduction

Sequencing throughput has improved rapidly in the last several years [1] while cost has continued to drop [2]. RNA sequencing (RNA-seq) [3, 4], a common use of sequencing, involves isolating and sequencing mRNA from biological samples. RNA-seq has become a standard tool for studying gene expression due to its ability to detect novel transcriptional activity in a largely unbiased manner without relying on previously defined gene sequence. Indeed, the Sequence Read Archive contains data for over 170,000 RNA-seq samples, including over 45,000 from human samples [5]. Large-scale projects like GTEx [6] and TCGA [7] are generating RNA-seq data on thousands of samples across normal and malignant tissues derived from hundreds to thousands of individuals.

The goal of an RNA-seq experiment is often to characterize expression across all samples and identify features associated with outcomes of interest. The first step, read alignment, determines where sequencing reads originated with respect to the reference genome or annotated transcriptome. Unlike read alignment for DNA sequencing reads, RNA-seq alignment should be splice-aware to accommodate introns that have been spliced out of the mature mRNA transcripts, creating an exon-exon junction. While RNA-seq data can be generated on hundreds or thousands of samples, the alignment process has up to this point largely been performed on each sample separately [8–24].

We introduce Rail-RNA, a splice-aware, annotation-agnostic read aligner designed to analyze many RNA-seq samples at once. Rail-RNA makes maximal use of data from many samples by (a) borrowing strength across replicates to achieve accurate detection of exon-exon junctions, even at low coverage, and (b) avoiding effort spent aligning sequences that are redundant either within or across samples. Furthermore, Rail-RNA can be run on a computer cluster at the user’s home institution or on a cluster rented from a commercial cloud computing provider at a modest cost per sample. Cloud services rent out standardized units of hardware and software, enabling other Rail-RNA users to easily reproduce large-scale analyses performed in other labs. Rail-RNA’s output is compatible with downstream tools for isoform assembly, quantitation, and count- and region-based differential expression.

We demonstrate Rail-RNA is more accurate than other tools and is increasingly accurate as more samples are added. We show Rail-RNA is less susceptible to biases affecting other tools; specifically, (a) Rail-RNA has substantially higher recall of alignments across low-coverage exon-exon junctions and (b) Rail-RNA is accurate without a gene annotation, avoiding annotation bias resulting from potentially incomplete [25] or incorrect transcript annotations. We run Rail-RNA on 667 paired-end lymphoblastoid cell line (LCL) RNA-seq samples from the GEUVADIS study [26], obtaining results in 15 hours and 47 minutes at a cost of US$0.91^1^ per sample. This is a fraction of per-sample sequencing costs, which are $20 or more [27]. We illustrate the usability of Rail-RNA’s output by performing a region-based differential expression analysis of the GEUVADIS dataset. Our analysis identifies 285,695 expressed regions, including 19,649 that map to intergenic sequence. We further show that intergenic and annotated regions show similar patterns of variation across populations and with respect to known technical confounders.

Altogether, Rail-RNA is a significant step in the direction of usable software that quickly, reproducibly, and accurately analyzes large numbers of RNA-seq samples at once.

## 2 Results

The GEUVADIS consortium performed mRNA sequencing of 465 lymphoblastoid cell line samples derived from CEPH (CEU), Finnish (FIN), British (GBR), Toscani (TSI) and Yoruba (YRI) populations of the 1000 Genomes Project [26], giving 667 paired-end samples. Per-sample sequencing depth is summarized in Supplementary Material, Section 5.1. For information on reproducing all our results, including software versions, see Supplementary Material, Section 5.2.

### 2.1 Design principles of Rail-RNA

Rail-RNA follows the MapReduce programming model, and is structured as an alternating sequence of aggregate steps and compute steps. Aggregate steps group and order data in preparation for future compute steps. Compute steps run concurrently on ordered groups of data. In this framework, it is natural to aggregate data across samples, i.e. to bring together related data so decisions can be informed by all samples at once. This affords greater accuracy and efficiency than if samples are analyzed separately. Rail-RNA aggregates across samples at multiple points in the pipeline to increase accuracy (borrowing strength for junction calling) and scalability (through elimination of redundant alignment work).

Rail-RNA can be run in elastic, parallel, or single-computer mode. In single-computer mode, Rail-RNA runs on multiple processors on a single computer. In parallel mode, Rail-RNA runs on any of the variety of cluster setups supported by IPython parallel [28]. These include Sun Grid Engine (SGE), Message Passing Interface (MPI), and StarCluster.

Elastic mode uses Hadoop [29], an implementation of MapReduce [30]. In elastic mode, Rail-RNA is run using the Amazon Web Services (AWS) Elastic MapReduce (EMR) service. EMR is specifically for running Hadoop applications on computer clusters rented on demand from Amazon’s Elastic Compute Cloud (EC2). Amazon Simple Storage Service (S3) stores intermediate data and output. There are important advantages and disadvantages to commercial cloud computing services like AWS [31, 32]. One advantage is that it facilitates reproducibility: one researcher can reproduce the same hardware and software setup used by another. Another advantage for Rail-RNA, which stores all intermediate and final results in S3, is that there is no risk of exhausting the cluster’s disk space even for datasets with many samples. Without these facilities, making scalable software that runs easily on large numbers of RNA-seq samples in different laboratories is quite challenging.

Rail-RNA supports both paired-end and unpaired RNA-seq samples. It supports input data consisting of reads of various lengths, and can detect exon-exon junctions involving exons as short as 9 bp by default. Rail-RNA uses Bowtie 2 [33] to align reads, including reads spanning exon-exon junctions, so it is naturally both indel-aware (with affine gap penalty) and base quality-aware. Rail-RNA is also deterministic; the same input data and parameters will yield the same outputs regardless of number of processors and computers used and regardless of whether it runs in elastic, parallel, or single-computer mode.

### 2.2 Scalability

We randomly selected 112 paired-end samples from the GEUVADIS study, with 16 coming from each of the 7 sequencing laboratories. We also randomly selected subsets of 28 and 56 samples from the 112 to illustrate Rail-RNA’s scalability on EMR (Supplementary Material, Section 5.3).

We use the term “instance” to refer to a computer (or virtualized fraction of one) that can be rented from EC2. An instance can have multiple processing cores. For experiments described in this section: (1) we used c3.2xlarge EC2 spot instances, each with eight processing cores. See Supplementary Material, Section 5.4 for details on this instance type and Supplementary Material, Section 5.5 for details on spot instances; (2) we used Amazon’s Simple Storage Service (S3) to store inputs, intermediate results, and final results; (3) we performed all experiments and stored all S3 data in the EU region (eu-west-1). See Supplementary Material, Section 5.6 for details on cost measurements. For every experiment in this paper, we measure cost by totaling the bid price for the EC2 spot instances (here, $0.11 per instance per hour using spot marketplace) and the Elastic MapReduce surcharges (here, $0.105 per instance per hour). On the one hand, this estimate can be low since it does not account for costs incurred by S3 and related services. On the other, the estimate can be high since the actual price paid depends on the spot market price, which is lower than the bid price.

The 112 selected GEUVADIS samples spanned 2.4 terabytes of compressed FASTQ data. We preprocessed each subset of 28, 56, and 112 samples of the 112 using different Elastic MapRe-duce clusters, each spanning 21 c3.2xlarge instances. Each cluster downloaded source data from a European Nucleotide Archive FTP server [34, 35]. Preprocessing 28 samples took 1 hour and 3 minutes, costing $9.03, preprocessing 56 samples took 1 hour and 14 minutes, costing $9.03, and preprocessing 112 samples took 2 hours and 14 minutes, costing $13.54.

**Figure 1:**
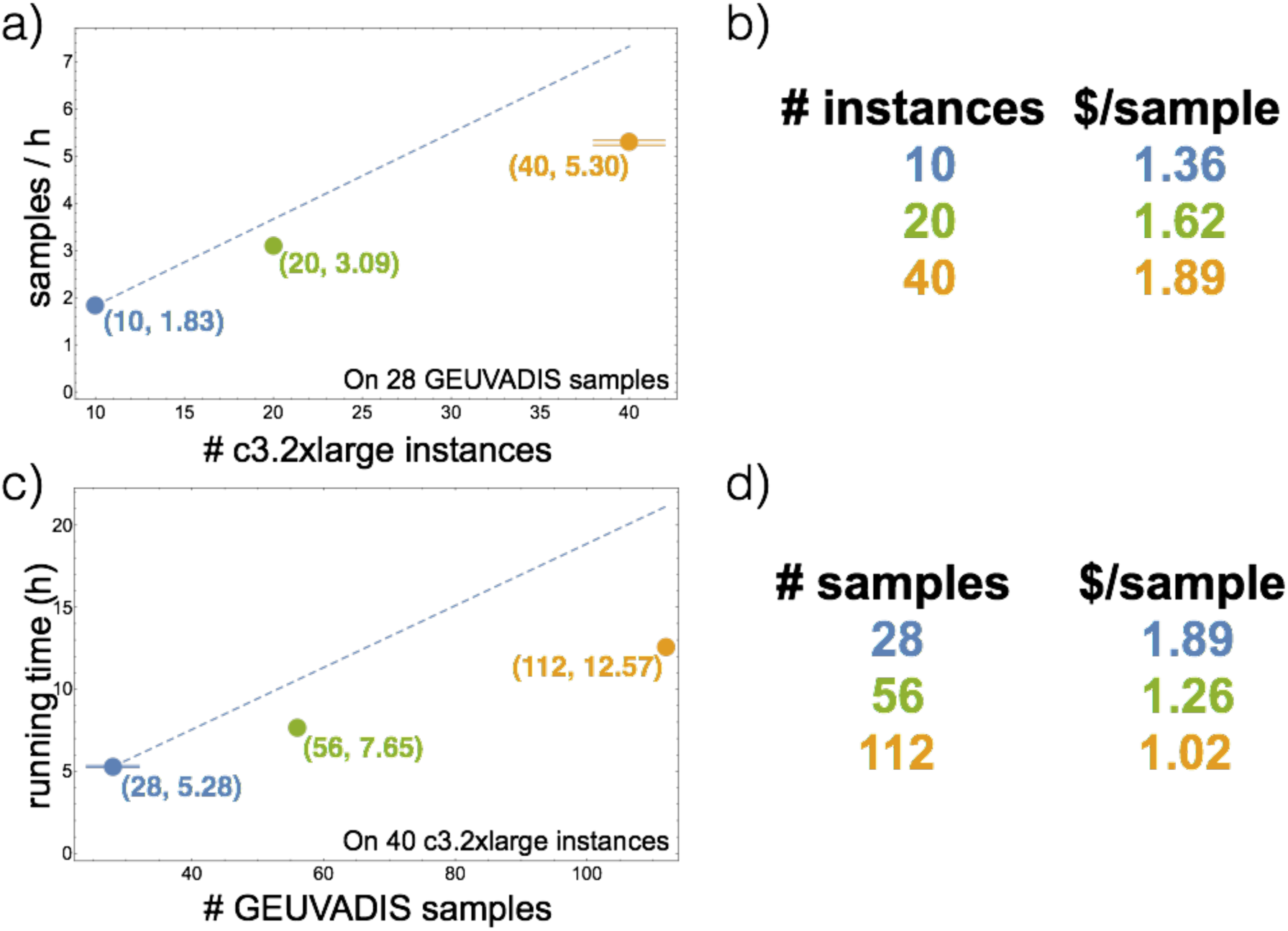
a) depicts scaling with respect to cluster size. Horizontal axis is the number of instances in the cluster. Vertical axis is throughput measured as number of samples analyzed per hour. The dashed line illustrates ideal linear scaling, extrapolated from the 10-instance result. b) is a table of per-sample costs for each experiment in a). c) plots Rail-RNA’s running time on 28, 56, and 112 paired-end GEUVADIS samples. The dashed line represents linear scaling, extrapolating that it takes twice as long to analyze approximately twice as much data. Rail-RNA achieves better-than-linear scaling with respect to number of samples as reflected in the table of costs in d): cost per sample decreases as number of samples analyzed increases. Per-sample costs in b) and d) reported here do not include the cost of preprocessing, which depends on download speed. These costs are reported in the main text. The master instance is also not included in the cluster sizes quoted here.

We ran Rail-RNA several times to assess scalability, which we measured with respect to both the number of instances in the cluster and the number of input samples. To measure scalability with respect to cluster size, Rail-RNA was run on a random subset of 28 of the 112 samples using EMR clusters of 10, 20, and 40 c3.2xlarge core instances (Figure 1a). (Each EMR cluster has an extra instance called the master instance that coordinates cluster activity; we exclude it from instance counts here because it contributes no workers.) Each of the 10and 20-instance experiments was run exactly once, while the 40-instance experiment was run three times to measure wall-clock time variability. The orange dot representing the experiment on 40 instances is at the mean number of samples analyzed per hour, while the horizontal lines above and below represent the minimum and maximum values. The dashed blue line shows an ideal linear extrapolation from the 10-instance data point. This illustrates how throughput would increase if doubling the number of instances also doubled throughput. Rail-RNA’s scaling falls short of ideal linear scaling, but this is expected due to various sources of overhead including communication and load imbalance. Importantly, Rail-RNA’s throughput continues to grow as the number of instances increases to 40, indicating Rail-RNA can make effective use of hundreds of processors at once.

To measure scalability with respect to number of input samples, Rail-RNA was run on 28, 56, and 112 samples using a cluster of 40 instances (Figure 1b). The 40-instance experiment with 28 samples (blue dot and lines) reports the same results as in 1a, but now in terms of running time. The dashed blue line extrapolates linear scaling from the 28-sample data point, assuming doubling the number of samples doubles running time. The 56and 112-sample points fall well below the line, indicating Rail-RNA achieves better-than-linear scaling of running time with respect to number of samples analyzed. Rail-RNA gets more efficient as more samples are analyzed in part because it identifies and eliminates redundant alignment work within and across samples (Supplementary Material, Section 5.8). Analyzing many samples together is particularly beneficial, with cost per sample dropping from $1.89 for 28 samples to $1.02 for 112 samples (Figure 1d). In an experiment described in Section 2.4, per-sample cost was reduced to $0.91 per sample.

A breakdown of the time taken by the steps in Rail-RNA’s pipeline is provided in Supplementary Material, Section 5.7.

### 2.3 Accuracy

We simulated 112 RNA-seq samples with 40 million 76-bp paired-end reads each using Flux Simulator [36]. Typically, Flux assigns expression levels to transcripts according to a modified Zipf’s law. Instead, we used FPKM expression levels from the set of 112 randomly selected paired-end samples studied in Section 2.2; they are taken from the GEUVADIS study [26] and are available at [37]. See Supplementary Material, Section 5.9 for simulation details.

We compared Rail-RNA’s accuracy to that of TopHat 2 v2.1.0 [10], Subjunc from v1.4.6p-v4 of the Subread package [38], STAR v2.4.2a [11], and HISAT v0.1.6-beta [39]. We ran TopHat 2 with (“Tophat 2 ann”) and without (“Tophat 2 no ann”) the Gencode v12 annotation [40] provided. We ran Subjunc using the default values of its command-line parameters. We ran STAR in three ways: in one pass (“STAR 1 pass”); in one pass with exon-exon junctions from Gencode v12 provided (“STAR 1 pass ann”); and in two passes (“STAR 2 pass”). We similarly ran HISAT in three ways: in one pass (“HISAT 1 pass”); in one pass with exon-exon junctions from Gencode v12 provided (“HISAT 1 pass ann”); and in two passes (“HISAT 2 pass”). One-pass methods (“STAR 1 pass,” “STAR 1 pass ann,” “HISAT 1 pass,” and “HISAT 1 pass ann”) align reads and call exon-exon junctions in one step, whereas two-pass methods (all other protocols) additionally perform a second step that realigns reads in light of exon-exon junction calls from the first. Supplementary Material, Section 5.10 describes the protocols in detail.

When an alignment program is run with annotation (“Tophat 2 ann,” “STAR 1 pass ann,” and “HISAT 1 pass ann”), we provide the same annotation from which Flux Simulator simulated the reads. That is, the provided annotation consists of a superset of the actual transcripts present. Consequently, protocols where the annotation is provided have an artificial advantage.

Rail-RNA was run in three ways:

1. **On a single sample (“Rail single”).** Rail-RNA uses reads from the given sample to identify exon-exon junctions. Like in two-pass protocols, reads are then realigned to transcript fragments to handle the case where a read overlaps an exon-exon junction near its 5’ or 3’ end.
2. **On 112 samples with exon-exon junction filter (“Rail all”).** After initial alignment, Rail-RNA compiles a list of exon-exon junctions found in any sample. The list is then filtered; an exon-exon junction passes the filter if (a) it appears in at least 5% of the input samples, or (b) it is overlapped by at least 5 reads in any sample. The filtered exon-exon junction list is used to build the set of transcript fragments to which each sample is then realigned.
3. **On 112 samples without exon-exon junction filter (“Rail all NF”)**. Identical to “Rail all” but with no exon-exon junction filter. This is not a recommended protocol; we include it only to show the filter’s effect.

In none of the three modes does Rail-RNA use a gene annotation: Rail-RNA is consistently annotation-agnostic.

We consider two accuracy measures in the main text: (1) overlap accuracy, measuring precision and recall of overlap events. Each event is an instance where the primary alignment of a read overlaps an exon-exon junction; (2) exon-exon junction accuracy, measuring precision of exon-exon junctions called by a given aligner and recall of the set of exon-exon junctions within a sample or across samples. We also compute F-score, the harmonic mean of precision and recall. Supplementary Material, Section 5.11 formally defines these measures as well as a measure of overall mapping accuracy.

The “Rail all” and “Rail all NF” protocols were run on all 112 simulations. Other protocols were run on each of the 20 samples randomly selected from the 112. We emphasize that even though protocols labeled “Rail all” analyze all 112 samples at once, we evaluate their output alignments and calls only for the sample of 20. Figure 2 displays overlap and exon-exon junction accuracy measurements. Figure 2a summarizes the accuracy of each tool across the 20 samples.

As illustrated in Figure 2a, Rail-RNA has the highest overlap F-score of the protocols tested, including those using a gene annotation. Rail-RNA’s overlap precision is comparable to the most precise protocol (“Subjunc” in this case) and its recall is comparable to the highest of any protocol (“STAR 1 pass ann”). Further, analyzing many samples at once (“Rail all”) achieves greater Fscore compared to analyzing one (“Rail single”). This is more pronounced for exon-exon junction accuracy than for overlap accuracy since borrowing strength across replicates is particularly effective at detecting low-coverage junctions (Supplementary Material, Section 5.13). Mean exon-exon junction recall increases from 0.880 to 0.939 (Supplementary Material, Section 5.12), which adds about 10,000 true positives.

Figure 2b demonstrates the efficacy of the exon-exon junction filter in the “Rail all” protocol. Precision/recall are defined similarly as for a single sample, but pooled across all samples. That is, recall is the fraction of exon-exon junctions with at least one simulated spanning read in at least one sample that Rail-RNA detects in at least one sample. The improvement in precision when moving from the unfiltered (0.673) to the filtered (0.964) exon-exon junction list shows that the filter removes a large fraction of incorrect exon-exon junction calls because they are supported in only a few samples. Further, the distribution of filtered exon-exon junctions with certain donor/acceptor motifs matches the expected distribution (GT-AG: 96.5%, GC-AG: 2.6%, AT-AC: 1.0%) much more closely than the same distribution for the unfiltered exon-exon junctions.

**Figure 2:**
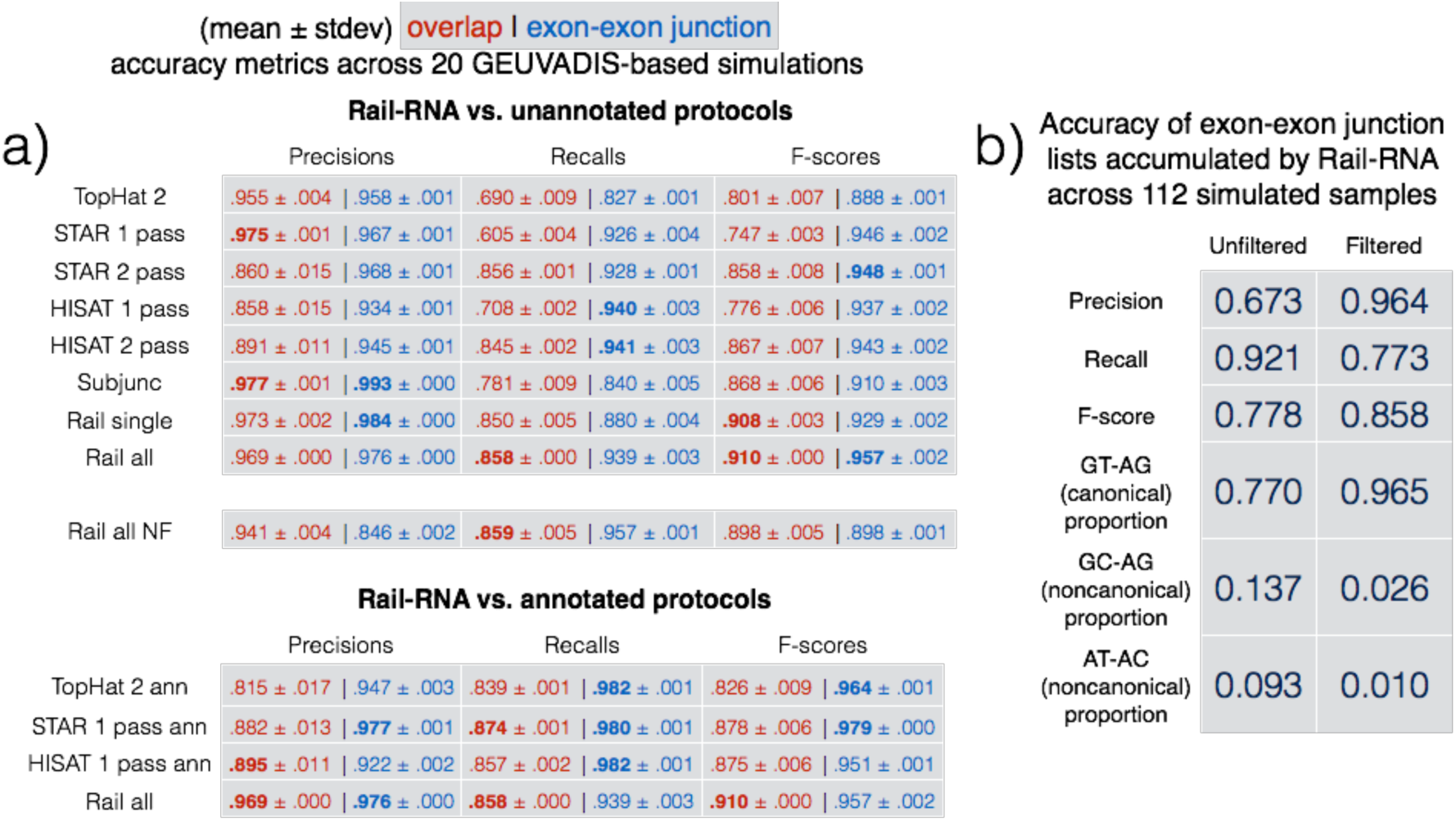
(a) Means and standard deviations of accuracy measures of various alignment protocols across 20 simulated samples whose FPKMs mimic those of 20 GEUVADIS samples. “Rail all” protocols involve running Rail-RNA on all 112 simulated samples. Overlap accuracy is shown in red, exon-exon junction accuracy in blue. Some tools are run with (top) and without (bottom) a provided gene annotation. Note that Rail-RNA never uses an annotation. For Rail-RNA vs. unannotated protocols, the best two results in each column are in boldface, and for Rail-RNA vs. annotated protocols, the best result in each column is in boldface. (b) Rail-RNA’s accuracy on two sets of exon-exon junctions found across all 112 simulated samples: one before and one after application of the exon-exon junction filter.

Rail-RNA’s protocols also tie “STAR 1 pass” in achieving the highest mean F-scores in an accuracy comparison that considers all alignments, including those falling entirely within exons; see Supplementary Material, Section 5.12.

To compare Rail-RNA’s speed on a single computer to that of other tools, we additionally measured wall-clock times of all single-sample alignment protocols for the GEUVADIS sample with SRA accession number ERR205018 on 8 and 16 processing cores in Supplementary Material, Section 5.14.

### 2.4 Analysis of GEUVADIS RNA-seq samples

We demonstrate the utility of Rail-RNA’s outputs by performing a novel analysis of 667 paired-end GEUVADIS RNA-seq samples from lymphoblastoid cell lines (LCLs) [26]. Starting from FASTQ inputs, Rail-RNA produced bigWig files [41] encoding genomic coverage; per-sample BED files recording exon-exon junctions, insertions and deletions; and sorted, indexed BAM [42] files recording the alignments. We downloaded and preprocessed all the GEUVADIS data using a cluster of 21 c3.2xlarge Amazon EC2 instances in 7 hours and 29 minutes, costing a total of $36.12. Rail-RNA then ran on 61 c3.8xlarge Amazon EC2 instances (see 5.4) spanning 1,952 processors. The run completed in 15 hours and 47 minutes and cost $605.12 ($0.91 per sample). Including preprocessing, the cost per sample was $0.96.

**Figure 3:**
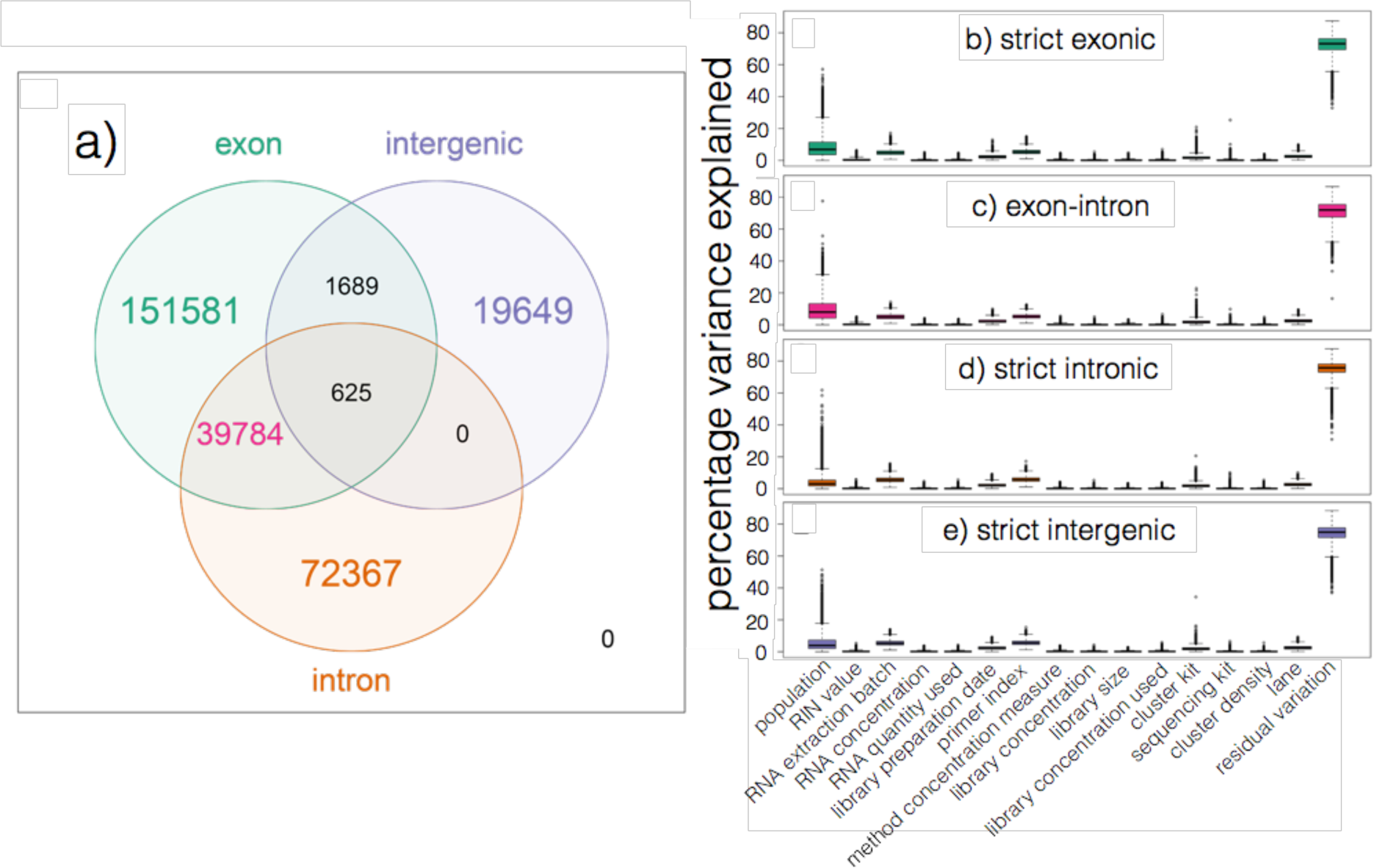
(a) Number of expressed regions (ERs) from the 667 GEUVADIS samples overlapping known exons, introns, and intergenic sequence as determined from Ensembl v75. (b)-(e) Boxplots of percentage variance in expression explained by assorted variables (population and 14 technical variables) as well as residual “unexplained” variation restricted to (b) strictly exonic ERs, (c) ERs overlapping known exons and introns, (d) strictly intronic ERs, and (e) strictly intergenic ERs. Each expressed region corresponds to one point in each of the 15 *×* 4 boxplots.

We analyzed bigWig outputs using the `derfinder` Bioconductor package [43]. `derfinder` identified contiguous genomic stretches where average coverage across all samples was above a genomewide threshold (see Supplementary Material, Section 5.15). Adjacent bases above the threshold were grouped into “expressed regions” (ERs). We identified 285,695 ERs in this way; median ER length was 70 bp (interquartile range, IQR: 7-145). While gene annotation/structure is not used to identify ERs, the regions can be overlapped with a gene annotation to assess novel transcriptional activity, as shown in Figure 3a. Relative to Ensembl v75 [40], we found that 151,581 ERs (53.1%) were within exons (median length: 93 bp, IQR: 21-155) and another 38,784 (13.9%) overlapped both exonic and intronic sequence (median length: 132 bp, IQR: 76-269) perhaps representing alternative ends to known exons. We also found 72,367 regions (25.3%) contained within intronic sequence (median length: 9 bp, IQR: 2-37) and another 19,649 regions (6.9%) within intergenic sequence (median length: 15 bp, IQR: 3-71). These intronic and intergenic ERs could represent novel (polyadenlyated, since the data was generated using polyA+ prep) non-coding RNA.

We also reproduced the variance component modeling illustrated in Figure 3 of Hoen et al 2013 [44]. At each of the 285,695 ERs, we modeled log_2_-adjusted expression levels as a function of many largely technical variables related to the RNA (RIN value, RNA concentration, and RNA quantity), the sequencing libraries (library preparation date, library primer index for each sample, method then target and actual quantities of library concentrations, and library size) and the sequencing itself (cluster kit, sequencing kit, cluster density and sequencing lane). We further included the ethnicity of each sample as a variable because it appeared to explain moderate variability in expression levels at many ERs. Lastly, we calculated the amount of residual variability not explained by any of the variables we considered. Interestingly, we found little difference in the amounts of variability explained by each of the variables studied when we stratified the ERs by the annotation categories described above (Figure 3b-e), suggesting that the technical factors affect expression of previously unannotated (i.e. intronic and intergenic) features in a similar manner as they do annotated gene structures.

## 3 Methods

For each read, Rail-RNA seeks the (possibly spliced) alignment maximizing the alignment score, which measures similarity between the read and reference. Bowtie 2’s local-mode [45] scoring function is used: matched bases add a bonus while mismatched bases and gaps incur a penalty. Gap penalties are affine. Ties for best alignment score are broken as follows. Each alignment *i* among the highest-scoring alignments overlaps some number *n*(*i*) of exon-exon junctions, where *n*(*i*) = 0 for an alignment wholly contained in a single exon. If the smallest *n*(*i*) among the highest-scoring alignments is attained by exactly one alignment, that alignment is chosen. If the smallest *n*(*i*) is attained by more than one alignment, the tie is broken in a weighted random fashion where alignments overlapping high-coverage junctions are preferred to alignments overlapping lowcoverage junctions (Supplementary Material, Section 5.16).

Methods used in the statistical analysis of the GEUVADIS dataset are described in Supplementary Material, Section 5.15. Steps of the Rail-RNA pipeline are described in the following sections. The term “workers” refers to computer processes under Rail-RNA’s control. Usually many workers operate simultaneously, distributed across several computers. Each step writes either intermediate or final output data. Depending on the outputs requested by the user, some steps may be omitted. Details of how Rail-RNA is implemented in various distributed environments are given in Supplementary Material, Section 5.17. Figure 4 illustrates how Rail-RNA eliminates redundant alignment work.

### 3.1 Preprocess reads

Initially, the user provides a manifest file containing a URL pointer to each input FASTQ file. Two URLs are specified for each paired-end sample, one for a single-end sample. URLs point to the local filesystem, the web, or on Amazon’s Simple Storage Service (S3). In this step, input reads are downloaded as necessary and preprocessed into a format that facilitates parallelism (Supplementary Material, Section 5.18).

### 3.2 Align reads to genome

Duplicate reads and readlets both within and across samples lead to redundant alignment work. To eliminate redundant work, Rail-RNA groups duplicate reads so that a worker operates on all reads having the same nucleotide sequence at once. Afterwards, two passes of alignment are performed using Bowtie 2. In the first, each unique read sequence is aligned to the genome. If there is exactly one highest-scoring alignment and it has no gaps, mismatches or soft-clipped bases, all reads with the same nucleotide sequence are assigned that alignment. If the alignment is not perfect or if there is more than one highest-scoring alignment, all reads with the same nucleotide sequence are run through a second pass of Bowtie 2 to ensure that quality sequences are taken into consideration when scoring alignments or ties are broken. Some read sequences with imperfect alignments are divided into short overlapping substrings called readlets. These sequences are searched for whether they overlap exon-exon junctions in a later step. See Supplementary Material, Section 5.19 for further details.

### 3.3 Align readlets to genome

Rail-RNA groups duplicate readlets so that a worker operates on all readlets across samples with the same nucleotide sequence at once. Each distinct readlet sequence is aligned to the genome eith Bowtie [46] exactly once, eliminating redundant alignment work. Rail-RNA searches for several possible alignments, up to 30 by default using command-line parameters `-a -m 30`.

### 3.4 Detect exon-exon junctions using readlet alignments

Rail-RNA uses correlation clustering and maximum clique finding to detect exon-exon junctions spanned by each distinct read sequence, as detailed in Supplementary Material, Section 5.20. The algorithm is reminiscent of the seed-and-vote strategy of Subread/subjunc [38], and we note similarities and differences in Supplementary Material, Section 5.21. The step outputs exon-exon junctions and the number of reads covering each junction in each sample.

### 3.5 Filter exon-exon junctions violating confidence criteria

In simulations we observed that junctions detected in only a small fraction of samples tend to be false positives (Figure 2b). Consequently, the global list of exon-exon junctions is quickly dominated by false positives as the number of samples increases. To keep precision high, Rail-RNA borrows strength across samples to remove junctions not meeting one of these criteria:

1. The exon-exon junction appears in at least *K*% of samples.
2. The exon-exon junction is covered by at least *J* reads in at least one sample.

*K* = *J* = 5 by default, but both are configurable.

### 3.6 Enumerate intron configurations

Rail-RNA enumerates the ways that multiple exon-exon junctions detected on the same strand in the same sample can be overlapped by a read segment **s**(*readlet_config_size*) spanning *readlet_config_size* bases. One possible way that a read or readlet can overlap multiple exon-exon junctions is called an “intron configuration.” Intron configurations for readlets are obtained and output as described in Supplementary Material, Section 5.23.

**Figure 4:**
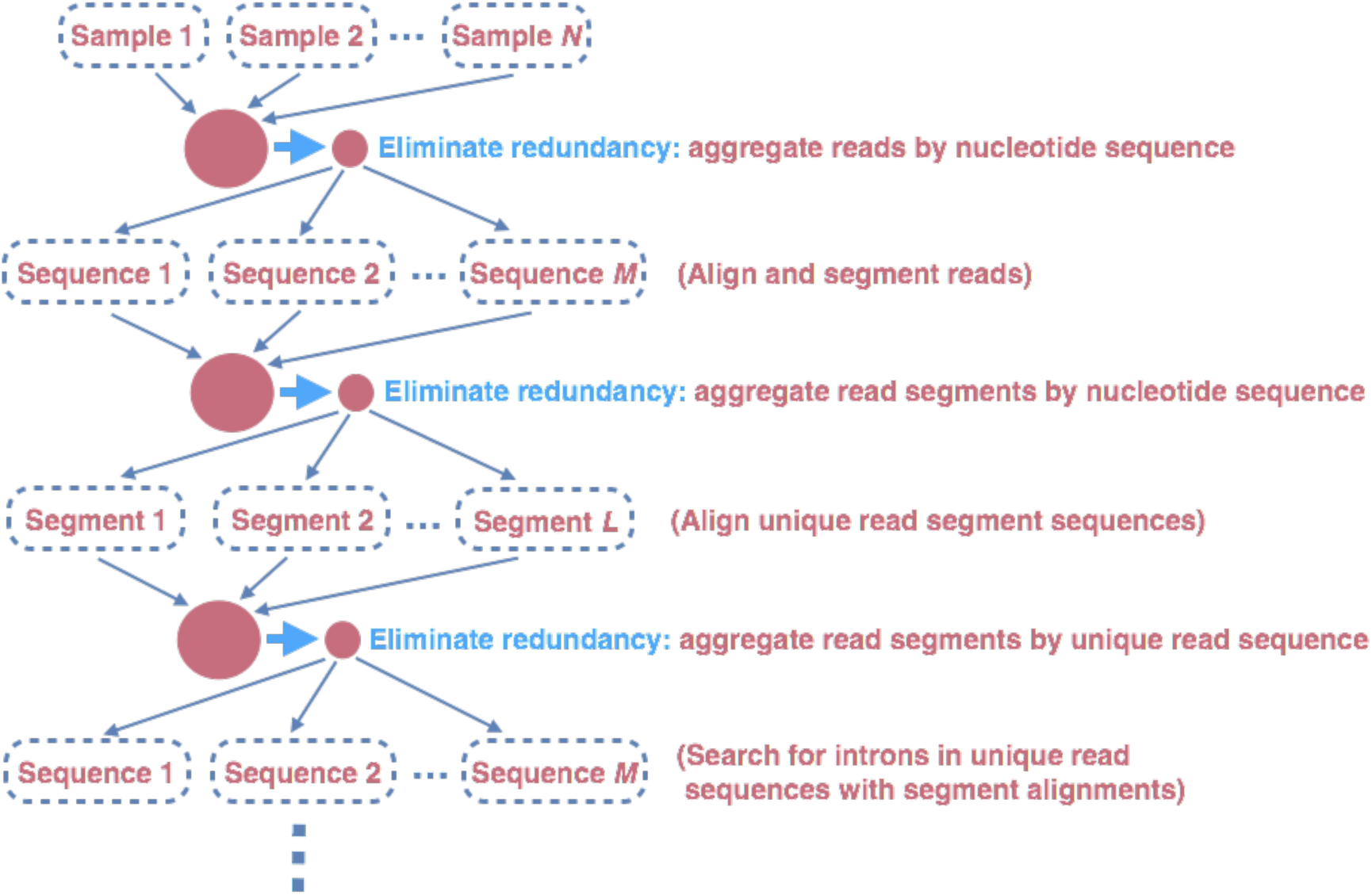
Rail-RNA eliminates redundant alignment work by first aggregating reads by nucleotide sequence and, where appropriate, aligning only unique read sequences to the genome. Read sequences that do not align perfectly to the genome are divided into overlapping segments, and after these segments are aggregated, each unique segment is aligned to the genome exactly once. Rail-RNA then gathers all segment alignments belonging to a read sequence and searches for exon-exon junctions within each unique read sequence.

### 3.7 Retrieve and index isofrags

Each worker operates on an intron configuration at a time, concatenating the exonic bases surrounding its introns to form a transcript fragment of size *readlet_config_size*. This is termed an “isofrag.” Care is taken to avoid including intronic bases in isofrags (Supplementary Material, Section 5.24). Subsequently, a single worker uses `bowtie2-build` to build a single Bowtie 2 index for all enumerated isofrags. Later, Bowtie 2 uses the index to realign reads in the next step.

### 3.8 Finalize combinations of exon-exon junctions overlapped by read sequences

Read sequences that failed to align perfectly in the first step are aligned to isofrags using Bowtie 2 in `local` mode with a minimum score threshold of 48 by default. Local alignment is used since indexed sequences are of length *readlet_config_size*, shorter than the read length. Rail-RNA runs Bowtie 2 with the parameter `-k 30` by default so that many alignments are reported per read sequence. From these alignments Rail-RNA derives a list of exon-exon junctions the read could possibly overlap. A graph algorithm enumerates the combinations of exon-exon junctions the read sequence might simultaneously overlap; see Supplementary Material, Section 5.25 for details.

### 3.9 Realign reads

Read sequences that failed to align perfectly in the first step are realigned to a set *S* of transcript fragments. Each transcript fragment in *S* overlaps a different combination of exon-exon junctions found in the previous step. All the exon-exon junction combinations found for the read’s nucleotide sequence are spanned by a *subset* of *S*. Moreover, several distinct read sequences may overlap transcript fragments in *S*. A given worker performs realignment as follows.

1. Transcript fragments in *S* are recorded and indexed with `bowtie2-build`.
2. Reads are realigned to the new index using Bowtie 2 in `--local` mode. These are reads that are in the same index bin referenced in Supplementary Material, Section 5.19.

### 3.10 Collect and compare read alignments

Bowtie 2 alignments of reads accumulated in previous steps, except for those that aligned perfectly in the “Align reads to genome” step (Section 3.2), are collected here and partitioned by read name. A worker operates on all alignments of a given read at once. For each read, if there is exactly one highest-scoring alignment for that read, it is chosen as the primary alignment. Otherwise, Rail-RNA attempts to break the tie by selecting the alignment spanning the fewest exon-exon junctions. If there is still a tie, it is broken by a random draw weighted by the number of uniquely aligned reads supporting each exon-exon junction, as described by (3) in Supplementary Material, Section 5.16.

### 3.11 Write BAMs recording alignments

By default, all primary alignments, including perfect alignments from the “Align reads to genome” step, are output here. The user may disable this output or instead specify the `-k X` parameter to ask Rail-RNA to output up to X highest-scoring alignments per read. Alignments are written as sorted, indexed BAM files. By default, one BAM file is output per sample per chromosome. In elastic mode, all BAM files and their indexes are uploaded to S3. Tools such as the UCSC genome browser [47] allow users to visualize portions of BAM files without having to download them first.

### 3.12 Compile coverage vectors and write bigWigs

By default, Rail-RNA records vectors encoding depth of coverage at each position in the reference genome. The user may disable this output. Two bigWig files are produced per sample: one records coverage of the genome by reads for which each has exactly one highest-scoring alignment, and the other records coverage of the genome by primary alignments. bigWig files encoding mean and median coverage of the genome across samples are also written. The contributions of each sample to the mean and median are normalized by the number of mapped reads in the sample. In elastic mode, bigWig files are uploaded to S3. For example, the analysis in Section 2.4 was performed on bigWig files on S3. Further, tools such as the UCSC genome browser [47] allow users to visualize portions of bigWig files without having to download them first. These methods are detailed in Supplementary Material, Section 5.26.

### 3.13 Write exon-exon junctions and indels

Rail-RNA writes a set of TSV files. Each file contains a table with rows corresponding to samples and columns to distinct features. The (*i, j*)th element is the number of reads in the *i*th sample containing the *j*th feature. Three TSVs are written per sample, one where features are insertions, one for deletions, and one for exon-exon junctions. Also, three BED files are written per sample: one with exon-exon junctions, one with insertions, and one with deletions. These are formatted identically to TopHat’s analogous output [48]. In elastic mode, these files are uploaded to S3, where they can be analyzed as soon as Rail-RNA completes. The user may disable any outputs of this step.

## 4 Discussion

Designing software that is both usable and able to run on many samples at a time is technically challenging. Rail-RNA demonstrates that this comes with unique and substantial advantages. Rail-RNA achieves better-than-linear growth in throughput (and consequently reduction in cost) per sample as the number of samples grows. By using information from all samples when analyzing data from a given sample, Rail-RNA achieves superior accuracy, with its accuracy advantage growing as samples are added. Rail-RNA also substantially resolves two biases affecting other tools: bias against low-coverage junctions and annotation bias. Rail-RNA results obtained by one investigator can be reliably reproduced by another since Rail-RNA computer clusters rented from commercial cloud services have standardized hardware and software.

We demonstrated Rail-RNA by re-analyzing a 667-sample dataset in 15 hours and 40 minutes at a per-sample cost of $0.91, far lower than sequencing cost [27]. We analyzed region-level differential expression by simply passing Rail-RNA’s bigWig outputs to standard downstream tools (e.g. the `derfinder` Bioconductor packages) for further analysis. Users can reproduce this analysis using resources rented form a commercial cloud provider, without having to downloading any large datasets. Altogether, this as an important step in the direction of usable software that quickly, reproducibly, and accurately analyzes large numbers of RNA-seq samples at once.

We used the simple “5 reads or 5% of samples” filter for calling exon-exon junctions to avoid overfitting to our simulations. Further investigation is needed to find how the filter should be adjusted to optimize accuracy, and how to account for other factors like sequencing depth or variability in junction profiles between samples. The filter also contributes to an “*N* + 1” problem: if Rail-RNA is used to analyze *N* replicates, but an *N* + 1^*th*^ replicate arrives later, it is difficult to analyze just the *N* + 1^*th*^ and produce the same output as if all *N* + 1 had been analyzed together. These are areas for future work, discussed further in Supplementary Material, Section 5.22 and Supplementary Material, Section 5.27.

## Acknowledgments

We thank the anonymous reviewers for their constructive comments. AN, LCT, JTL, and BL were supported by NIH/NIGMS grant 1R01GM105705 to JTL. AN was supported by a seed grant from the Institute for Data Intensive Engineering and Science (IDIES) at Johns Hopkins University to BL and JTL. LCT was supported by Consejo Nacional de Ciencia y Tecnología México 351535. BL and JP were supported by a Sloan Research Fellowship to BL. JAH was supported by Fundación UNAM. BL was supported by National Science Foundation grant IIS-1349906. Experiments using Amazon Web Services were supported by two AWS in Education grants to BL.

## 5. Supplementary material

### 5.1 GEUVADIS samples

Figure 5 depicts the distribution of counts of reads (counting ends of paired-end reads separately) across the 667 paired-end GEUVADIS RNA-seq samples. The median number of reads in a sample is 52, 609, 404 with median absolute deviation 12, 553, 184. Details on experimental and sequencing protocols are provided in the supplementary note of [44]. Each GEUVADIS sample has either 75or 76-bp reads.

**Figure 5:**
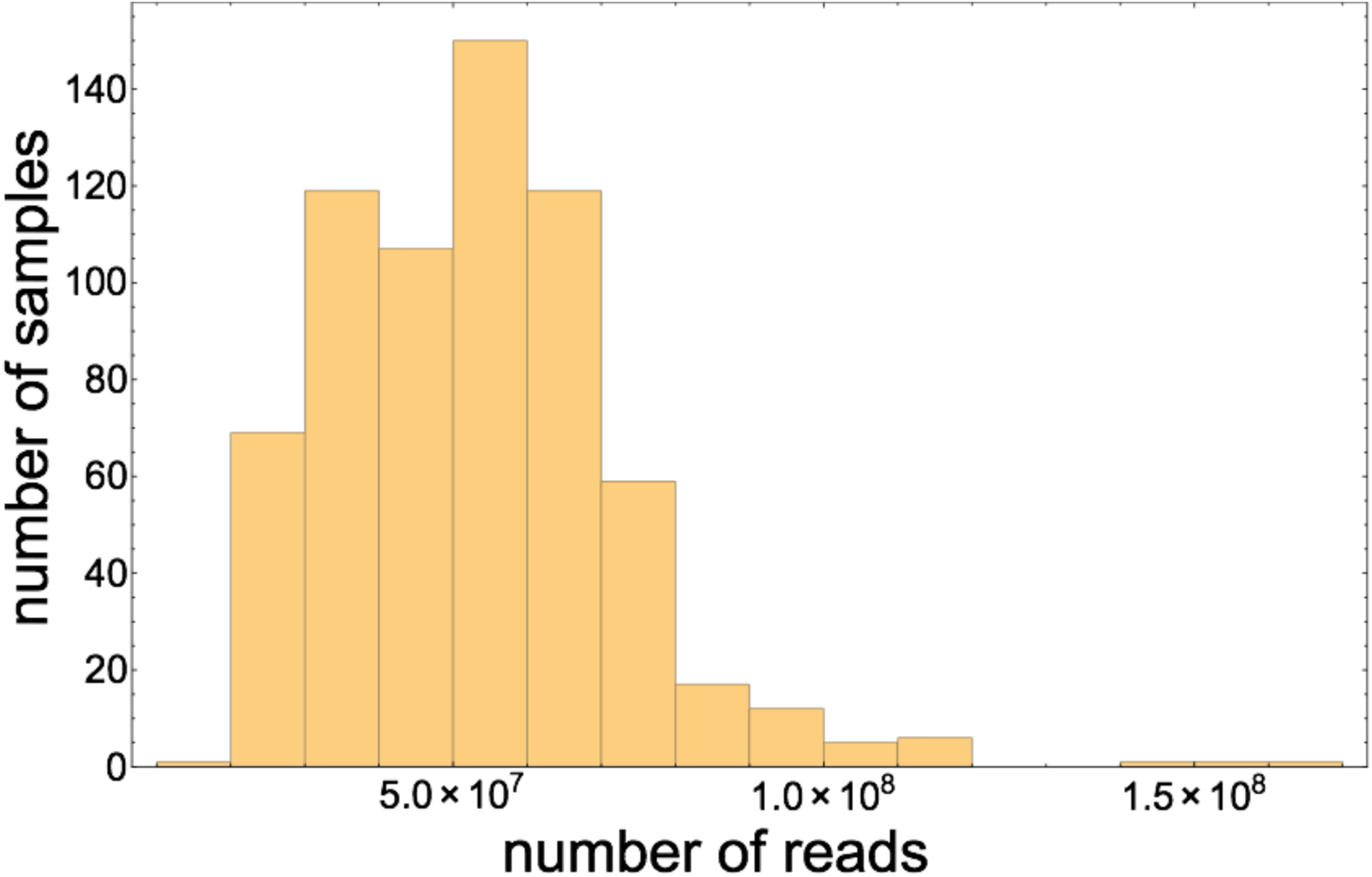
Histogram of counts of reads (not read pairs) in all 667 paired-end GEUVADIS samples.

### 5.2 Reproducing results

All scripts used to obtain the results in this paper are available at https://github.com/nellore/rail/tree/master/eval. Instructions on how to reproduce our results may be found at https://github.com/nellore/rail/blob/master/eval/README.md.

Rail-RNA v0.1.9b reproduces results in this paper and has several dependencies. In the experiments we conducted, Rail-RNA wrapped Bowtie 1 v1.1.1, Bowtie 2 v2.2.5, PyPy v2.5, and SAMTools v1.2. Version 2.1.0 of TopHat 2 was used and, like Rail-RNA, it wrapped Bowtie 2 v2.2.5. Version 2.4.2a of STAR was used, and version 0.1.6-beta of HISAT was used. Subjunc is a tool in the Subread package, and version 1.4.6-p4 of the Subread package was used. Flux Simulator 1.2.1 was used to obtain simulated samples.

### 5.3 GEUVADIS subsets

The 28 samples spanned 1,638,479,586 reads, the 56 samples spanned 3,127,479,220 reads, and the 112 samples spanned 6,402,672,808 reads. When we discuss Rail-RNA’s scaling with respect to number of samples, we implicitly assume that doubling the number of samples roughly doubles the number of reads. This is approximately true for the GEUVADIS subsets we consider.

### 5.4 Amazon Elastic Compute Cloud instance specifications

All experiments in this paper were conducted on either c3.2xlarge instances or c3.8xlarge instances. A c3.2xlarge instance is a virtualized computer powered by Intel Xeon E5-2680 v2 (Ivy Bridge) 64-bit processors that provides 15 GB of RAM, 320 GB of solid-state storage, and 8 2.8-Ghz processing cores. A c3.8xlarge instance is a virtualized computer powered by Intel Xeon E5-2680 v2 (Ivy Bridge) processors that provides 60 GB of RAM, 640 GB of solid-state storage, and 32 2.8-Ghz processing cores [49].

### 5.5 Spot instances

Spot instances permit bidding for EC2 computing capacity. If a bid exceeds the market price, which varies continuously according to supply and demand, requested computing capacity is made available to the user. A risk of using spot instances is that the computing capacity may be lost if market price rises to exceed the original bid price. Market prices vary from region (with an Amazon Web Services data center) to region. In July 2015, when our experiments were run, the market price in the EU region was under $0.09 per hour per c3.2xlarge instance. By contrast, reserving a c3.2xlarge instance in July 2015 in the EU region on demand, which ensures no job failure due to fluctuations in market price, cost $0.42 per hour in July 2015.

### 5.6 Measuring Amazon Web Services costs

For every experiment in this paper, we measure cost by totaling the bid price for the EC2 spot instances ($0.11 per c3.2xlarge instance per hour and $0.35 per c3.8xlarge instance using the spot marketplace) and the Elastic MapReduce surcharges ($0.105 per c3.2xlarge instance per hour and $0.27 per c3.8xlarge instance per hour). On the one hand, this estimate can be low since it does not account for costs incurred by S3 and related services. On the other, the estimate can be high since the actual price paid depends on the spot market price, which is lower than the bid price.

There are no onetime costs associated with setting up an Amazon Web Services account or with launching an Elastic MapReduce job.

The Rail-RNA “Preprocess reads” step (Section 3.1) is a necessary first step to any analysis of data with Rail-RNA. To calculate end-to-end time and cost of an experiment, i.e. the time and cost of the experiment starting from raw data situated in an archive, one must sum the time and cost of the preprocessing step and the rest of the Rail-RNA pipelines. These costs are all reported in the main text.

### 5.7 Breakdown of step times for scaling experiments

**Table 1:** Breakdown of step times for scaling experiments. Each row is a step, each column is an experiment. Experiments are labeled as the ordered pair (number of GEUVADIS samples analyzed, number of c3.2xlarge instances used), except that 28-DUP refers to the experiment labeled as 28DUP in Section 5.8. WCT is wall-clock time including provisioning, bootstrapping, and Hadoop setup, while WCT-B is wall-clock time excluding provisioning, bootstrapping, and Hadoop setup. Units are minutes except where marked “s” for seconds.

|  | (28, 10) | (28, 20) | (28-DUP, 40) | (28, 40) | (28, 40) | (28, 40) | (56, 40) | (112, 40) |
| --- | --- | --- | --- | --- | --- | --- | --- | --- |
| Provisioning, bootstrapping, and Hadoop setup | 40 | 54 | 38 | 48 | 56 | 50 | 45 | 44 |
| Align reads and segment them into readlets | 207 | 124 | 86 | 74 | 75 | 75 | 117 | 223 |
| Align readlets to genome | 34 | 18 | 6 | 9 | 9 | 9 | 14 | 24 |
| Detect junctions using readlet alignments | 87 | 52 | 12 | 23 | 23 | 22 | 40 | 70 |
| Filter junctions that violate confidence criteria | 1 | 58 s | 56 s | 56 s | 54 s | 56 s | 58 s | 1 |
| Enumerate possible junction overlaps on readlets | 1 | 58 s | 56 s | 56 s | 54 s | 56 s | 58 s | 1 |
| Get isofragments for index construction | 54 s | 54 s | 54 s | 52 s | 52 s | 54 s | 54 s | 54 s |
| Build isofragment index | 1 | 1 | 1 | 2 | 1 | 1 | 1 | 1 |
| Finalize junction overlap combinations on reads | 92 | 50 | 14 | 27 | 27 | 28 | 29 | 52 |
| Get transcriptome elements for read realignment | 14 | 10 | 3 | 4 | 4 | 4 | 4 | 7 |
| Align reads to isofragments | 260 | 133 | 62 | 70 | 68 | 69 | 117 | 162 |
| Collect and compare read alignments | 76 | 39 | 23 | 19 | 19 | 19 | 34 | 67 |
| Associate spliced alignments with junction coverages | 8 | 4 | 2 | 2 | 2 | 3 | 4 | 7 |
| Finalize primary alignments of spliced reads | 6 | 3 | 2 | 2 | 2 | 2 | 2 | 4 |
| Write BAMs with alignments by sample | 41 | 23 | 13 | 12 | 12 | 12 | 22 | 41 |
| Write mapped read counts | 40 s | 36 s | 38 s | 36 s | 36 s | 38 s | 36 s | 38 s |
| Merge exon differentials at same genomic positions | 16 | 9 | 5 | 5 | 5 | 5 | 8 | 17 |
| Compile sample coverages from exon differentials | 9 | 5 | 3 | 3 | 3 | 2 | 4 | 9 |
| Write bigWigs with exome coverage by sample | 15 | 10 | 9 | 8 | 8 | 7 | 10 | 16 |
| Aggregate junctions/indels | 5 | 3 | 2 | 2 | 2 | 2 | 3 | 5 |
| Write normalization factors/junctions/indels | 1 | 1 | 1 | 1 | 1 | 1 | 1 | 56 s |
| Write BEDs with junctions/indels by sample | 1 | 1 | 1 | 1 | 56 s | 1 | 56 s | 1 |
| WCT | 917 | 543 | 287 | 315 | 321 | 315 | 459 | 754 |
| WCT-B | 877 | 489 | 249 | 267 | 265 | 265 | 414 | 710 |

### 5.8 Measuring the effect of eliminating redundant alignment work

Wall-clock times in Figure 1 include time taken to provision Elastic MapReduce clusters, install required software and Bowtie indexes on the clusters using bootstrap scripts, and set up Hadoop. This extra time is largely independent of cluster size across experiments and should properly be subtracted from wall-clock times (WCT) when studying the scaling of strictly the Rail-RNA pipeline. The row WCT-B of Table 1 excludes the time taken to set up clusters from the wall-clock time. Rail-RNA’s mean WCT-B value for 28 samples on 41 c3.2xlarge instances is about 266 minutes. Extrapolating this value linearly to 56 and 112 samples gives 532 minutes and 1,064 minutes, respectively, exceeding the actual WCT-B values of, respectively, 489 and 877 minutes. So Rail-RNA exhibits better-than-linear scaling with respect to number of samples when the time cost of provisioning and setting up clusters is excluded.

That said, it is difficult to attribute better-than-linear scaling directly (or mostly) to elimination of redundant alignment work. There may be other difficult-to-measure sources of overhead that lead to better-than linear scaling. For example, any source of overhead that takes constant time with respect to the number of input samples (e.g. Java Virtual Machine startup time) could have a similar effect.

Nevertheless, we can provide strong evidence that Rail-RNA invests less time in aligning datasets composed of more “similar” samples, i.e. samples with greater sequence redundancy. We selected 14 of the 28 GEUVADIS samples on which we ran our experiments measuring Rail-RNA’s scalability with respect to cluster size in Section 2.2 and duplicated each sample, giving it a new sample name. In the resulting set of 28 samples—call it 28-DUP—each read in one sample was *guaranteed* to have at least one exactly identical read in another sample, increasing redundant sequence in the dataset. 28-DUP spanned 1,725,465,520 reads (not read pairs) in total, while the original set of 28 GEUVADIS samples (28-ORIG) spanned 1,638,479,586 reads. The WCT for running Rail-RNA on 28-DUP on 41 c3.2xlarge instances was 287 minutes, while the WCT-B was 249 minutes (Table 1). Contrast these values with the mean WCT and WCT-B for 28-ORIG on 41 c3.2xlarge instances: 317 minutes and 266 minutes, respectively. So despite including 86,985,934 more reads (about 1.5 GEUVADIS samples’ worth) than 28-ORIG, 28-DUP took 17 fewer minutes than the mean WCT-B for 28-ORIG to complete its run. Note that the range of variation in WCT-B when analyzing 28ORIG on 41 c3.2xlarge instances—2 minutes (see Table 1)—indicates that such variation is unlikely to account for the difference in analysis time between 28-ORIG and 28-DUP.

### 5.9 Simulated data

The 112 GEUVADIS samples on whose coverage distributions our simulated samples are based are the same as those studied in Section 2.2: all seven labs that pursued the study are represented, with sixteen samples from each lab. FPKMs were previously computed in the GEUVADIS study [26] with Flux Capacitor [50] for transcripts from Gencode v12 [51] after an initial mapping to the GRCh37 (hg19) reference genome using GEM v1.349 [23]. For each sample, we interrupted the Flux Simulator pipeline after it had generated an expression profile and overwrote the appropriate intermediate file with the computed expression profile. More specifically, the final column of this intermediate file gives the number of molecules of each transcript to be simulated. We replaced the value for each transcript with 20 times its FPKM from Flux Capacitor. This gave a mean number of 15.5 million RNA molecules per simulation, ensuring adequate library yield in the remainder of the pipeline. After we resumed the pipeline, Flux Simulator simulated fragmentation, reverse transcription, and sequencing; for each sample, this yielded a FASTQ file with raw reads and a BED file specifying each read’s location in the genome. Flux Simulator’s built-in 76-base error model was used in the sequencing step. This model is biased towards generating more base substitutions near the low-quality (3’) ends of reads. Across 76-bp reads in all simulated samples, the mean rate of substitution errors is 2.50%. Define a read with a low-quality tail as a read for which each of the ten bases at its 3’ end has Phred score less than or equal to 15. 8.59% of all 76-bp reads across simulated samples have low-quality tails. Across 76-bp reads *without* low-quality tails in all simulated samples, the mean rate of substitution errors is 0.82%.

**Figure 6:**
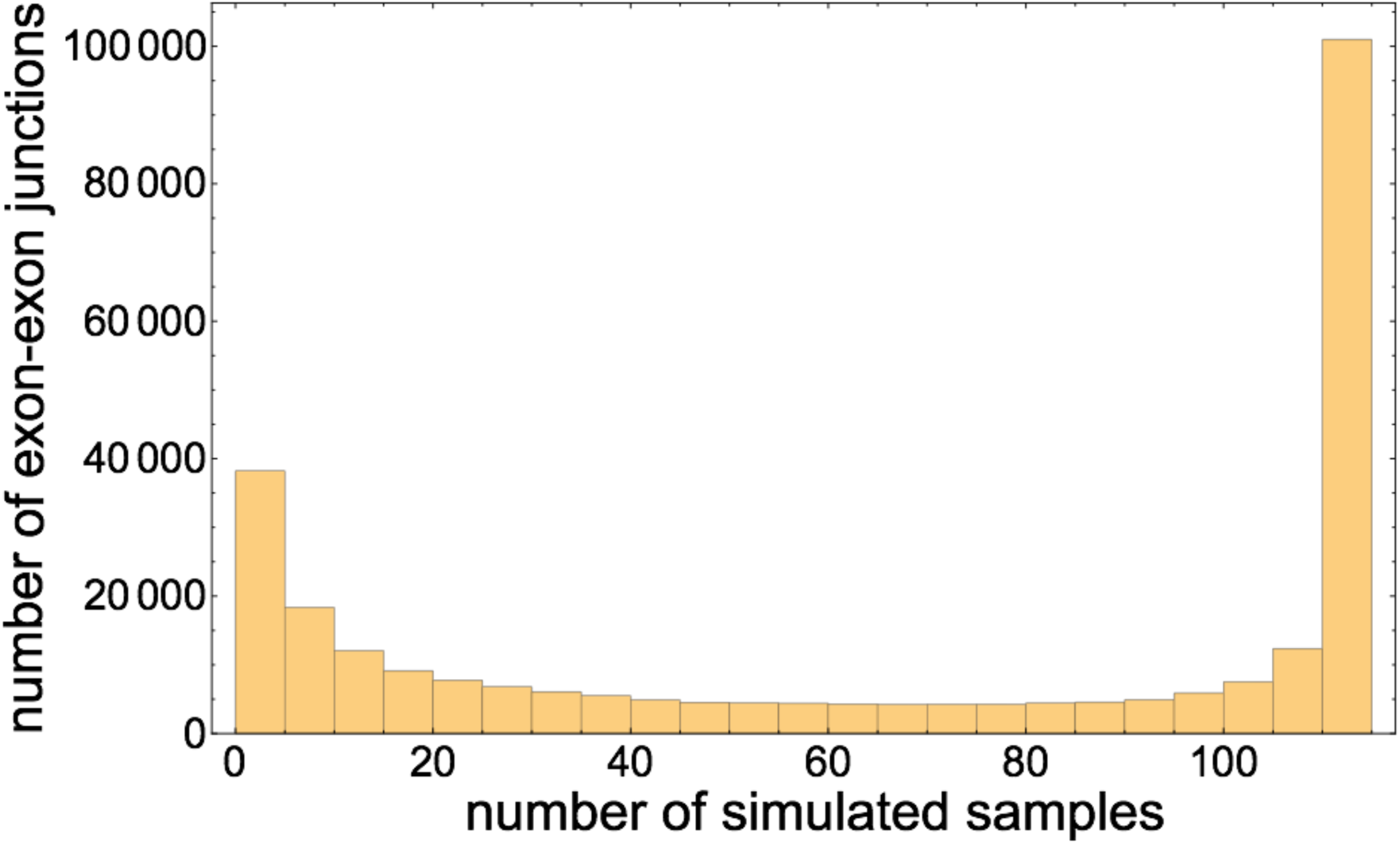
Histogram of the number of simulated samples out of 112 in which each exon-exon junction appears.

While the simulated samples were all generated from the same gene annotation, because each is designed to have a distinct coverage distribution, the exon-exon junction content of each sample is also distinct. Some junctions occur in many samples, while others occur in few. In particular, of 279, 681 distinct exon-exon junctions sampled across the 112 simulated samples, 5.86% occur in exactly one sample, and 15.28% occur in five or fewer samples. A core of 31.3% of exon-exon junctions occur in all 112 samples. Figure 6 illustrates that the distribution of the number of samples in which each exon-exon junction occurs is bimodal with peaks at tails.

The 112 simulated samples may be regenerated using the script https://github.com/buci/rail/tree/master/eval/generate_bioreps.py. Table 2 gives the URLs of the FASTQ files for the GEUVADIS sample on which each simulation was based. A sample number is given for a line if the corresponding simulated sample was one of the twenty randomly selected for alignment using all protocols considered in this paper and detailed in Section 5.10. The sample number coincides with the sample numbers referenced in the accuracy tables of Section 5.12.

**Table 2:**
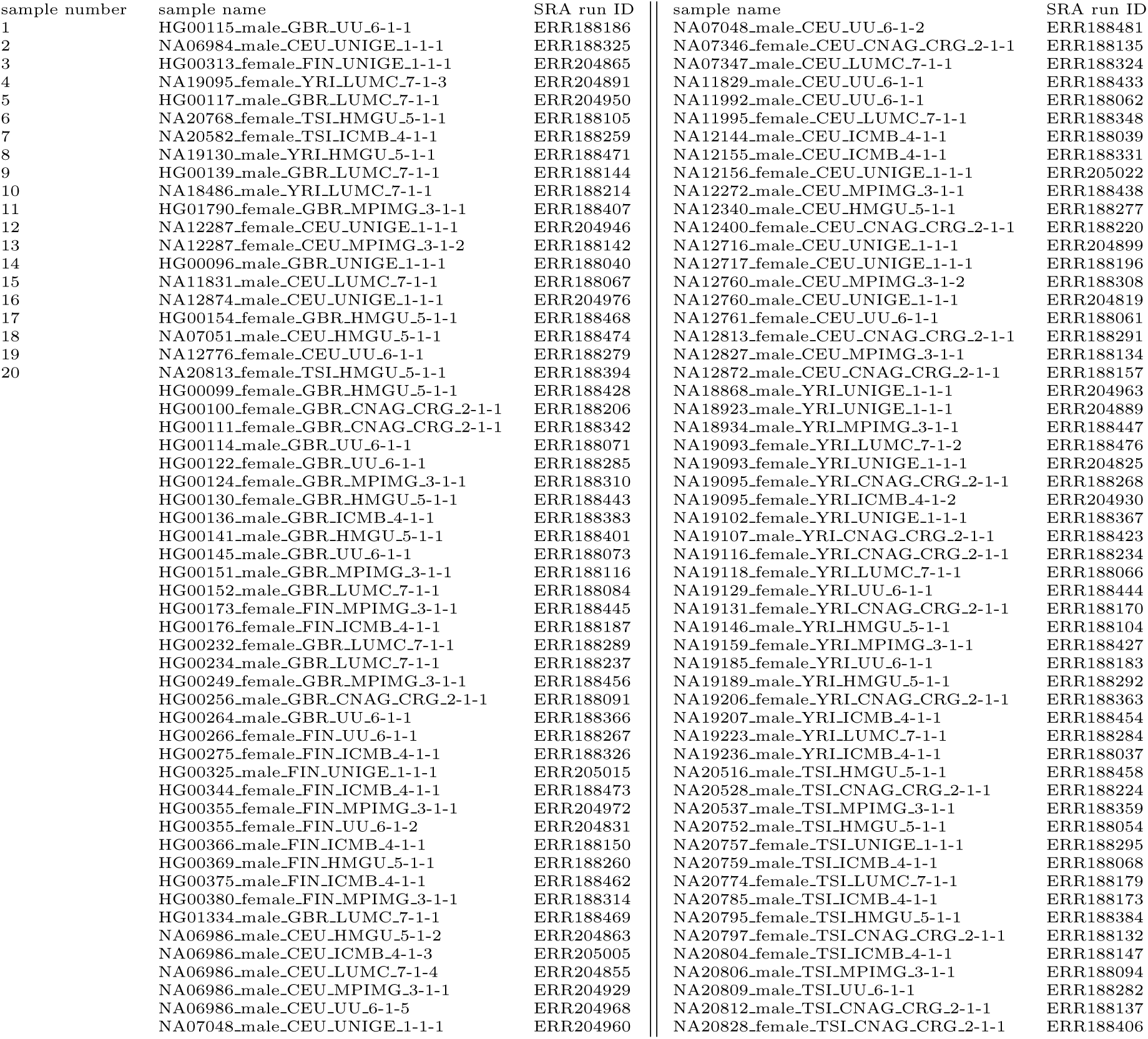
GEUVADIS samples on whose coverage distributions the 112 simulated RNA-seq samples from this paper are based. The 20 simulated samples referenced at the beginning of the table were randomly selected from the 112 for comparing Rail’s alignment protocols with alternative alignment protocols; their sample numbers correspond to the sample numbers from the accuracy tables in Section 5.12. Sample names are the names Rail-RNA used when aligning samples; they are in the format 

~~~
[Coriell sample ID]-[sex]-[HapMap population]-[GEUVADIS lab designator]-[lab ID (redundant)]-[biorep ID]-[techrep ID]
~~~

.

### 5.10 Alignment protocols

Commands executed for different alignment protocols may be generated by the script https://github.com/buci/rail/tree/master/eval/create_commands_for_all_sample_sims.py. Default command-line parameters were used for all protocols unless otherwise specified. In the “STAR 1 pass” protocol, STAR searches for exon-exon junctions in each individual read and does not realign reads. The “STAR 2 pass” protocol does not build a new index; rather, the protocol involves a single invocation of the STAR command with the parameters

`--twopass1readsN -1 --sjdbOverhang 75 --twopassMode Basic`, conforming to the protocol from the STAR manual [52]. We found empirically that an older two-pass protocol where

1. an index with the exonic bases surrounding exon-exon junctions compiled from the STAR 1 pass run is built. This index includes not just such transcript fragments, but also independently the full genome;
2. STAR is rerun to realign all reads to this index;
 gives almost exactly the same results. The older protocol is published in the supplement of the RGASP paper [53], which compares the accuracies of several spliced alignment protocols. Otherwise, the STAR protocols used here mirror those used for the RGASP paper.

In the “HISAT 1 pass” protocol, HISAT searches for exon-exon junctions in each read while also accumulating a list of exon-exon junctions on the fly for which to additionally search in reads. In “HISAT 2 pass” protocol,

1. HISAT was run in a mode where novel splice sites were output to a file using its `--novel-splicesite-outfile` parameter.
2. HISAT was rerun on all reads with as extra input the novel splice sites found in 1. using the `--novel-splicesite-infile` parameter.

### 5.11 Accuracy metrics

**Overlap accuracy**. Define a *true overlap* as the event that a read’s true alignment from simulation overlaps an exon-exon junction. Define a *positive overlap* as the event that a read’s primary alignment as reported by a given spliced alignment program overlaps an exon-exon junction. A *true positive* is a case where an exon-exon junction overlapped by the true alignment of a read is also overlapped by the reported primary alignment of the read. Precision is

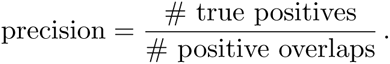

Recall is

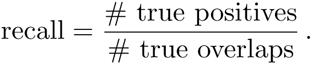

**Exon-exon junction accuracy**. Let *R* be the set of exon-exon junctions overlapped by at least one primary alignment reported by the aligner. Let *T* be the set of exon-exon junctions overlapped by at least one simulated read. Precision is

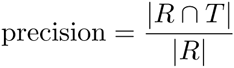

and recall is

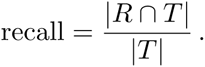

**Mapping accuracy**. Define a *true match* as the event that a given base of a read’s true alignment from simulation is aligned to a given genome position. A simulated read spanning *K* bases thus contributes *K* true matches and can be written as the ordered triple (**r**, *i, j*), where **r** is a read label, *i* is a position along the read, and *j* is the genome position to which *i* truly aligns. Define a *positive match* as the event that a given read base of a primary alignment reported by a given spliced alignment program is aligned to a given genome position. A positive match can similarly be represented as (**r**, *i, j*), where *j* is now the genome position to which the aligner aligned the read base. A *true positive* is a case where a true match equals a positive match.

Precision is

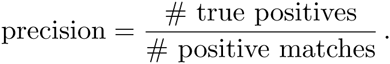

Recall is

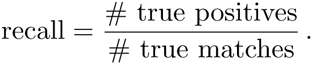

Note that this is a base-level measure of mapping accuracy that accounts for the fact that some aligners (i.e. Rail-RNA, STAR, and Subjunc) will produce a soft-clip alignment rather than leave a read unmapped. Since credit is given only for matched bases (and not soft-clipped bases), the contribution of soft-clipped alignments is not unduly inflated.

In this paper, the F-score is always the harmonic mean of precision and recall:

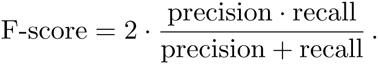

### 5.12 Accuracy tables

We present full tables corresponding to the accuracy results summarized in the main text. In each table, rows correspond to alignment protocols and columns to simulated samples—except for the last two columns; the next-to-last column records mean values across samples, and the final column records standard deviations across samples. Each element in each table has three numbers separated by commas: in order, they are always a precision, a recall, and an F-score. See Supplementary Material, Section 5.11 for descriptions of how precisions, recalls, and F-scores were computed for each table.

**Table 3:** Exon-exon junction accuracy for various alignment protocols described in Sections 2.3 and 5.10. Each table item is a tuple in the format “precision, recall, F-score”; these quantities are defined in Section 5.11. Sample numbers correspond to 20 randomly selected samples of the 112 samples we simulated, and they also appear in Table 2. Summary statistics are computed across the 20 samples. Note that the “Rail all” and “Rail all NF” protocols were each run on all 112 simulated samples to augment exon-exon junction calling.

| sample numbers | 1 | 2 | 3 | 4 | 5 |
| --- | --- | --- | --- | --- | --- |
| HISAT 1 pass | 0.934,0.940,0.937 | 0.934,0.937,0.936 | 0.933,0.944,0.938 | 0.935,0.941,0.938 | 0.934,0.939,0.936 |
| STAR 1 pass | 0.968,0.925,0.946 | 0.968,0.922,0.944 | 0.967,0.929,0.947 | 0.968,0.927,0.947 | 0.967,0.925,0.945 |
| HISAT 2 pass | 0.945,0.941,0.943 | 0.945,0.938,0.941 | 0.945,0.945,0.945 | 0.946,0.942,0.944 | 0.944,0.940,0.942 |
| Rail single | 0.984,0.880,0.929 | 0.984,0.875,0.926 | 0.983,0.883,0.930 | 0.984,0.882,0.930 | 0.984,0.878,0.928 |
| STAR 2 pass | 0.969,0.928,0.948 | 0.968,0.924,0.946 | 0.968,0.931,0.949 | 0.969,0.929,0.949 | 0.967,0.927,0.947 |
| Subjunc | 0.992,0.839,0.909 | 0.993,0.835,0.907 | 0.992,0.845,0.913 | 0.993,0.843,0.912 | 0.992,0.839,0.909 |
| TopHat 2 | 0.957,0.825,0.886 | 0.957,0.821,0.884 | 0.957,0.834,0.891 | 0.959,0.830,0.890 | 0.956,0.826,0.886 |
| HISAT 1 pass ann | 0.924,0.983,0.952 | 0.921,0.981,0.950 | 0.920,0.984,0.951 | 0.923,0.983,0.952 | 0.923,0.982,0.952 |
| STAR 1 pass ann | 0.977,0.980,0.979 | 0.977,0.979,0.978 | 0.977,0.981,0.979 | 0.977,0.981,0.979 | 0.977,0.980,0.978 |
| TopHat 2 ann | 0.947,0.981,0.964 | 0.946,0.980,0.963 | 0.945,0.983,0.964 | 0.946,0.982,0.963 | 0.948,0.982,0.964 |
| Rail all | 0.976,0.940,0.958 | 0.976,0.928,0.952 | 0.976,0.938,0.957 | 0.976,0.942,0.959 | 0.976,0.938,0.957 |
| Rail all NF | 0.845,0.956,0.897 | 0.850,0.954,0.899 | 0.848,0.957,0.899 | 0.843,0.958,0.897 | 0.847,0.955,0.898 |
| sample numbers | 6 | 7 | 8 | 9 | 10 |
| HISAT 1 pass | 0.934,0.939,0.937 | 0.935,0.939,0.937 | 0.933,0.937,0.935 | 0.934,0.939,0.937 | 0.934,0.938,0.936 |
| STAR 1 pass | 0.968,0.925,0.946 | 0.967,0.924,0.945 | 0.966,0.921,0.943 | 0.967,0.924,0.945 | 0.967,0.924,0.945 |
| HISAT 2 pass | 0.945,0.940,0.943 | 0.945,0.940,0.943 | 0.944,0.938,0.941 | 0.946,0.939,0.943 | 0.946,0.939,0.943 |
| Rail single | 0.984,0.878,0.928 | 0.984,0.878,0.928 | 0.984,0.875,0.926 | 0.984,0.879,0.928 | 0.984,0.877,0.928 |
| STAR 2 pass | 0.968,0.927,0.947 | 0.968,0.926,0.947 | 0.967,0.924,0.945 | 0.969,0.926,0.947 | 0.968,0.926,0.947 |
| Subjunc | 0.992,0.838,0.909 | 0.993,0.838,0.909 | 0.993,0.835,0.907 | 0.992,0.837,0.908 | 0.993,0.837,0.908 |
| TopHat 2 | 0.958,0.824,0.886 | 0.960,0.822,0.886 | 0.959,0.819,0.883 | 0.957,0.826,0.886 | 0.957,0.825,0.886 |
| HISAT 1 pass ann | 0.924,0.982,0.952 | 0.924,0.982,0.952 | 0.922,0.981,0.951 | 0.925,0.982,0.953 | 0.925,0.982,0.953 |
| STAR 1 pass ann | 0.978,0.980,0.979 | 0.977,0.980,0.979 | 0.976,0.979,0.978 | 0.978,0.980,0.979 | 0.978,0.980,0.979 |
| TopHat 2 ann | 0.948,0.982,0.965 | 0.949,0.980,0.964 | 0.948,0.980,0.964 | 0.950,0.982,0.965 | 0.950,0.982,0.965 |
| Rail all | 0.976,0.940,0.957 | 0.976,0.943,0.959 | 0.976,0.942,0.958 | 0.977,0.938,0.957 | 0.977,0.937,0.957 |
| Rail all NF | 0.847,0.956,0.898 | 0.844,0.958,0.897 | 0.843,0.957,0.896 | 0.850,0.955,0.900 | 0.850,0.955,0.900 |
| sample numbers | 11 | 12 | 13 | 14 | 15 |
| HISAT 1 pass | 0.935,0.945,0.940 | 0.934,0.946,0.940 | 0.934,0.943,0.938 | 0.934,0.941,0.937 | 0.934,0.939,0.936 |
| STAR 1 pass | 0.968,0.932,0.949 | 0.967,0.933,0.950 | 0.967,0.929,0.947 | 0.967,0.926,0.946 | 0.967,0.925,0.946 |
| HISAT 2 pass | 0.946,0.946,0.946 | 0.945,0.947,0.946 | 0.945,0.944,0.944 | 0.945,0.942,0.943 | 0.945,0.940,0.942 |
| Rail single | 0.984,0.887,0.933 | 0.984,0.889,0.934 | 0.984,0.883,0.931 | 0.984,0.881,0.929 | 0.984,0.878,0.928 |
| STAR 2 pass | 0.969,0.934,0.951 | 0.968,0.935,0.951 | 0.968,0.931,0.949 | 0.968,0.929,0.948 | 0.969,0.927,0.947 |
| Subjunc | 0.993,0.849,0.915 | 0.993,0.851,0.917 | 0.992,0.845,0.913 | 0.992,0.841,0.910 | 0.993,0.840,0.910 |
| TopHat 2 | 0.959,0.837,0.894 | 0.958,0.841,0.896 | 0.958,0.832,0.891 | 0.957,0.829,0.888 | 0.959,0.826,0.888 |
| HISAT 1 pass ann | 0.920,0.984,0.951 | 0.919,0.985,0.951 | 0.921,0.983,0.951 | 0.922,0.982,0.951 | 0.924,0.983,0.952 |
| STAR 1 pass ann | 0.976,0.982,0.979 | 0.976,0.983,0.979 | 0.976,0.981,0.979 | 0.977,0.980,0.979 | 0.977,0.980,0.978 |
| TopHat 2 ann | 0.943,0.983,0.963 | 0.942,0.984,0.963 | 0.945,0.983,0.964 | 0.947,0.982,0.964 | 0.949,0.981,0.965 |
| Rail all | 0.975,0.942,0.959 | 0.976,0.942,0.959 | 0.976,0.940,0.958 | 0.976,0.940,0.958 | 0.977,0.942,0.959 |
| Rail all NF | 0.844,0.959,0.898 | 0.847,0.959,0.899 | 0.847,0.957,0.898 | 0.846,0.956,0.898 | 0.848,0.957,0.899 |
| sample numbers | 16 | 17 | 18 | 19 | 20 |
| HISAT 1 pass | 0.934,0.946,0.940 | 0.934,0.938,0.936 | 0.936,0.937,0.936 | 0.934,0.937,0.935 | 0.933,0.937,0.935 |
| STAR 1 pass | 0.967,0.933,0.950 | 0.968,0.923,0.945 | 0.968,0.922,0.944 | 0.967,0.921,0.943 | 0.968,0.923,0.945 |
| HISAT 2 pass | 0.944,0.947,0.946 | 0.944,0.939,0.941 | 0.946,0.937,0.942 | 0.945,0.938,0.941 | 0.944,0.938,0.941 |
| Rail single | 0.984,0.888,0.934 | 0.984,0.877,0.927 | 0.985,0.876,0.927 | 0.984,0.874,0.926 | 0.983,0.877,0.927 |
| STAR 2 pass | 0.968,0.935,0.951 | 0.969,0.925,0.946 | 0.969,0.924,0.946 | 0.968,0.923,0.945 | 0.969,0.925,0.947 |
| Subjunc | 0.993,0.852,0.917 | 0.992,0.838,0.909 | 0.993,0.836,0.908 | 0.993,0.833,0.906 | 0.992,0.838,0.909 |
| TopHat 2 | 0.959,0.841,0.896 | 0.958,0.821,0.884 | 0.961,0.820,0.885 | 0.958,0.817,0.882 | 0.958,0.823,0.885 |
| HISAT 1 pass ann | 0.916,0.985,0.949 | 0.923,0.981,0.951 | 0.923,0.981,0.951 | 0.923,0.981,0.951 | 0.923,0.981,0.951 |
| STAR 1 pass ann | 0.975,0.982,0.979 | 0.978,0.979,0.978 | 0.977,0.979,0.978 | 0.977,0.979,0.978 | 0.977,0.979,0.978 |
| TopHat 2 ann | 0.940,0.984,0.961 | 0.949,0.980,0.964 | 0.947,0.980,0.963 | 0.949,0.980,0.964 | 0.948,0.980,0.964 |
| Rail all | 0.975,0.940,0.957 | 0.976,0.940,0.958 | 0.976,0.939,0.957 | 0.976,0.936,0.956 | 0.976,0.942,0.958 |
| Rail all NF | 0.845,0.959,0.898 | 0.847,0.956,0.898 | 0.848,0.956,0.899 | 0.847,0.955,0.898 | 0.844,0.957,0.897 |
| summary statistics | means | stddevs |  |  |  |
| HISAT 1 pass | 0.934,0.940,0.937 | 0.001,0.003,0.002 |  |  |  |
| STAR 1 pass | 0.967,0.926,0.946 | 0.001,0.004,0.002 |  |  |  |
| HISAT 2 pass | 0.945,0.941,0.943 | 0.001,0.003,0.002 |  |  |  |
| Rail single | 0.984,0.880,0.929 | 0.000,0.004,0.002 |  |  |  |
| STAR 2 pass | 0.968,0.928,0.948 | 0.000,0.004,0.002 |  |  |  |
| Subjunc | 0.993,0.840,0.910 | 0.000,0.005,0.003 |  |  |  |
| TopHat 2 | 0.958,0.827,0.888 | 0.001,0.007,0.004 |  |  |  |
| HISAT 1 pass ann | 0.922,0.982,0.951 | 0.002,0.001,0.001 |  |  |  |
| STAR 1 pass ann | 0.977,0.980,0.979 | 0.001,0.001,0.000 |  |  |  |
| TopHat 2 ann | 0.947,0.982,0.964 | 0.003,0.001,0.001 |  |  |  |
| Rail all | 0.976,0.939,0.957 | 0.000,0.003,0.002 |  |  |  |
| Rail all NF | 0.846,0.957,0.898 | 0.002,0.001,0.001 |  |  |  |

**Table 4:** Overlap accuracy for various alignment protocols described in Sections 2.3 and 5.10. Each table item is a tuple in the format “precision, recall, F-score”; these quantities are defined in Section 5.11. Sample numbers correspond to 20 randomly selected samples of the 112 samples we simulated, and they also appear in Table 2. Summary statistics are computed across the 20 samples. Note that the “Rail all” and “Rail all NF” protocols were each run on all 112 simulated samples to augment exon-exon junction calling.

| sample numbers | 1 | 2 | 3 | 4 | 5 |
| --- | --- | --- | --- | --- | --- |
| HISAT 1 pass | 0.857,0.709,0.776 | 0.854,0.709,0.775 | 0.876,0.707,0.782 | 0.875,0.709,0.783 | 0.862,0.706,0.776 |
| STAR 1 pass | 0.974,0.604,0.746 | 0.974,0.606,0.747 | 0.976,0.610,0.751 | 0.976,0.610,0.750 | 0.974,0.607,0.748 |
| HISAT 2 pass | 0.890,0.845,0.867 | 0.886,0.845,0.865 | 0.905,0.848,0.876 | 0.902,0.848,0.874 | 0.892,0.845,0.868 |
| Rail single | 0.973,0.849,0.907 | 0.970,0.849,0.905 | 0.974,0.857,0.912 | 0.976,0.857,0.913 | 0.973,0.853,0.909 |
| STAR 2 pass | 0.855,0.855,0.855 | 0.855,0.855,0.855 | 0.878,0.857,0.867 | 0.878,0.857,0.867 | 0.866,0.856,0.861 |
| Subjunc | 0.977,0.777,0.866 | 0.976,0.778,0.866 | 0.978,0.794,0.876 | 0.979,0.791,0.875 | 0.977,0.786,0.871 |
| TopHat 2 | 0.958,0.687,0.800 | 0.949,0.687,0.797 | 0.957,0.703,0.811 | 0.958,0.700,0.809 | 0.955,0.694,0.804 |
| HISAT 1 pass ann | 0.893,0.857,0.875 | 0.889,0.856,0.872 | 0.908,0.859,0.882 | 0.908,0.860,0.883 | 0.897,0.857,0.876 |
| STAR 1 pass ann | 0.875,0.874,0.875 | 0.873,0.873,0.873 | 0.896,0.874,0.885 | 0.900,0.875,0.887 | 0.890,0.873,0.881 |
| TopHat 2 ann | 0.806,0.838,0.822 | 0.802,0.837,0.819 | 0.833,0.838,0.836 | 0.839,0.840,0.839 | 0.824,0.837,0.831 |
| Rail all | 0.969,0.856,0.909 | 0.966,0.857,0.908 | 0.970,0.864,0.914 | 0.972,0.864,0.915 | 0.968,0.860,0.911 |
| Rail all NF | 0.940,0.857,0.897 | 0.937,0.858,0.896 | 0.944,0.866,0.903 | 0.947,0.865,0.904 | 0.940,0.861,0.899 |
| sample numbers | 6 | 7 | 8 | 9 | 10 |
| HISAT 1 pass | 0.856,0.707,0.774 | 0.840,0.711,0.770 | 0.840,0.710,0.770 | 0.889,0.707,0.788 | 0.873,0.706,0.781 |
| STAR 1 pass | 0.974,0.603,0.745 | 0.974,0.600,0.742 | 0.974,0.602,0.744 | 0.977,0.610,0.751 | 0.975,0.610,0.750 |
| HISAT 2 pass | 0.889,0.844,0.866 | 0.877,0.844,0.860 | 0.874,0.842,0.858 | 0.915,0.850,0.881 | 0.902,0.848,0.874 |
| Rail single | 0.973,0.848,0.906 | 0.972,0.843,0.903 | 0.972,0.845,0.904 | 0.975,0.858,0.913 | 0.975,0.856,0.912 |
| STAR 2 pass | 0.859,0.855,0.857 | 0.836,0.856,0.846 | 0.847,0.855,0.851 | 0.888,0.858,0.873 | 0.877,0.857,0.867 |
| Subjunc | 0.977,0.778,0.866 | 0.976,0.764,0.857 | 0.975,0.772,0.862 | 0.978,0.795,0.877 | 0.978,0.793,0.876 |
| TopHat 2 | 0.957,0.688,0.800 | 0.955,0.674,0.790 | 0.945,0.679,0.790 | 0.960,0.704,0.812 | 0.958,0.702,0.810 |
| HISAT 1 pass ann | 0.894,0.856,0.874 | 0.881,0.856,0.868 | 0.882,0.856,0.869 | 0.917,0.860,0.888 | 0.906,0.858,0.882 |
| STAR 1 pass ann | 0.879,0.874,0.877 | 0.861,0.875,0.868 | 0.870,0.875,0.873 | 0.904,0.874,0.889 | 0.897,0.875,0.886 |
| TopHat 2 ann | 0.811,0.839,0.825 | 0.788,0.840,0.813 | 0.798,0.839,0.818 | 0.846,0.838,0.842 | 0.834,0.839,0.836 |
| Rail all | 0.968,0.855,0.908 | 0.967,0.851,0.905 | 0.968,0.853,0.907 | 0.971,0.866,0.916 | 0.971,0.864,0.914 |
| Rail all NF | 0.939,0.857,0.896 | 0.936,0.852,0.892 | 0.936,0.854,0.893 | 0.949,0.867,0.906 | 0.945,0.865,0.903 |
| sample numbers | 11 | 12 | 13 | 14 | 15 |
| HISAT 1 pass | 0.859,0.711,0.778 | 0.873,0.707,0.782 | 0.859,0.707,0.776 | 0.869,0.709,0.781 | 0.858,0.711,0.777 |
| STAR 1 pass | 0.975,0.606,0.747 | 0.976,0.610,0.751 | 0.974,0.603,0.745 | 0.976,0.607,0.748 | 0.974,0.603,0.745 |
| HISAT 2 pass | 0.892,0.846,0.868 | 0.901,0.848,0.874 | 0.893,0.845,0.868 | 0.898,0.847,0.872 | 0.890,0.844,0.866 |
| Rail single | 0.975,0.851,0.909 | 0.974,0.858,0.912 | 0.973,0.849,0.907 | 0.973,0.852,0.909 | 0.973,0.847,0.906 |
| STAR 2 pass | 0.860,0.857,0.858 | 0.876,0.858,0.867 | 0.858,0.855,0.857 | 0.865,0.857,0.861 | 0.863,0.856,0.859 |
| Subjunc | 0.977,0.783,0.869 | 0.977,0.793,0.876 | 0.977,0.779,0.867 | 0.978,0.783,0.869 | 0.977,0.775,0.864 |
| TopHat 2 | 0.958,0.691,0.803 | 0.961,0.702,0.812 | 0.958,0.688,0.801 | 0.960,0.691,0.804 | 0.954,0.685,0.798 |
| HISAT 1 pass ann | 0.895,0.857,0.876 | 0.904,0.859,0.881 | 0.896,0.857,0.876 | 0.902,0.857,0.879 | 0.895,0.855,0.875 |
| STAR 1 pass ann | 0.879,0.875,0.877 | 0.894,0.874,0.884 | 0.880,0.874,0.877 | 0.883,0.874,0.878 | 0.883,0.875,0.879 |
| TopHat 2 ann | 0.813,0.840,0.826 | 0.834,0.838,0.836 | 0.814,0.839,0.826 | 0.818,0.838,0.828 | 0.817,0.839,0.828 |
| Rail all | 0.970,0.858,0.911 | 0.970,0.865,0.915 | 0.968,0.856,0.909 | 0.969,0.860,0.911 | 0.969,0.854,0.908 |
| Rail all NF | 0.942,0.859,0.899 | 0.945,0.866,0.904 | 0.942,0.857,0.898 | 0.942,0.861,0.899 | 0.940,0.856,0.896 |
| sample numbers | 16 | 17 | 18 | 19 | 20 |
| HISAT 1 pass | 0.864,0.708,0.778 | 0.839,0.707,0.767 | 0.839,0.706,0.767 | 0.842,0.709,0.770 | 0.842,0.711,0.771 |
| STAR 1 pass | 0.976,0.605,0.747 | 0.973,0.599,0.742 | 0.973,0.599,0.742 | 0.973,0.603,0.744 | 0.975,0.606,0.748 |
| HISAT 2 pass | 0.894,0.847,0.870 | 0.876,0.841,0.859 | 0.878,0.842,0.860 | 0.879,0.843,0.860 | 0.877,0.846,0.861 |
| Rail single | 0.976,0.852,0.909 | 0.971,0.844,0.903 | 0.972,0.844,0.903 | 0.971,0.846,0.904 | 0.973,0.853,0.909 |
| STAR 2 pass | 0.867,0.858,0.862 | 0.841,0.853,0.847 | 0.838,0.855,0.846 | 0.842,0.854,0.848 | 0.858,0.856,0.857 |
| Subjunc | 0.979,0.784,0.871 | 0.976,0.770,0.861 | 0.978,0.770,0.862 | 0.977,0.772,0.862 | 0.977,0.780,0.868 |
| TopHat 2 | 0.955,0.692,0.802 | 0.952,0.678,0.792 | 0.952,0.679,0.793 | 0.951,0.683,0.795 | 0.952,0.687,0.798 |
| HISAT 1 pass ann | 0.900,0.858,0.879 | 0.881,0.855,0.868 | 0.883,0.855,0.869 | 0.884,0.856,0.869 | 0.881,0.857,0.869 |
| STAR 1 pass ann | 0.886,0.875,0.880 | 0.865,0.874,0.869 | 0.863,0.875,0.869 | 0.864,0.873,0.869 | 0.889,0.875,0.882 |
| TopHat 2 ann | 0.823,0.840,0.831 | 0.793,0.838,0.815 | 0.791,0.840,0.815 | 0.791,0.838,0.814 | 0.817,0.839,0.828 |
| Rail all | 0.972,0.859,0.912 | 0.967,0.851,0.905 | 0.968,0.851,0.905 | 0.966,0.853,0.906 | 0.969,0.860,0.911 |
| Rail all NF | 0.944,0.860,0.900 | 0.935,0.852,0.892 | 0.933,0.852,0.891 | 0.934,0.854,0.893 | 0.941,0.861,0.899 |
| summary statistics | means | stdevs |  |  |  |
| HISAT 1 pass | 0.858,0.708,0.776 | 0.015,0.002,0.006 |  |  |  |
| STAR 1 pass | 0.975,0.605,0.747 | 0.001,0.004,0.003 |  |  |  |
| HISAT 2 pass | 0.891,0.845,0.867 | 0.011,0.002,0.007 |  |  |  |
| Rail single | 0.973,0.850,0.908 | 0.002,0.005,0.003 |  |  |  |
| STAR 2 pass | 0.860,0.856,0.858 | 0.015,0.001,0.008 |  |  |  |
| Subjunc | 0.977,0.781,0.868 | 0.001,0.009,0.006 |  |  |  |
| TopHat 2 | 0.955,0.690,0.801 | 0.004,0.009,0.007 |  |  |  |
| HISAT 1 pass ann | 0.895,0.857,0.875 | 0.011,0.002,0.006 |  |  |  |
| STAR 1 pass ann | 0.882,0.874,0.878 | 0.013,0.001,0.006 |  |  |  |
| TopHat 2 ann | 0.815,0.839,0.826 | 0.017,0.001,0.009 |  |  |  |
| Rail all | 0.969,0.858,0.910 | 0.002,0.005,0.003 |  |  |  |
| Rail all NF | 0.941,0.859,0.898 | 0.004,0.005,0.005 |  |  |  |

**Table 5:**
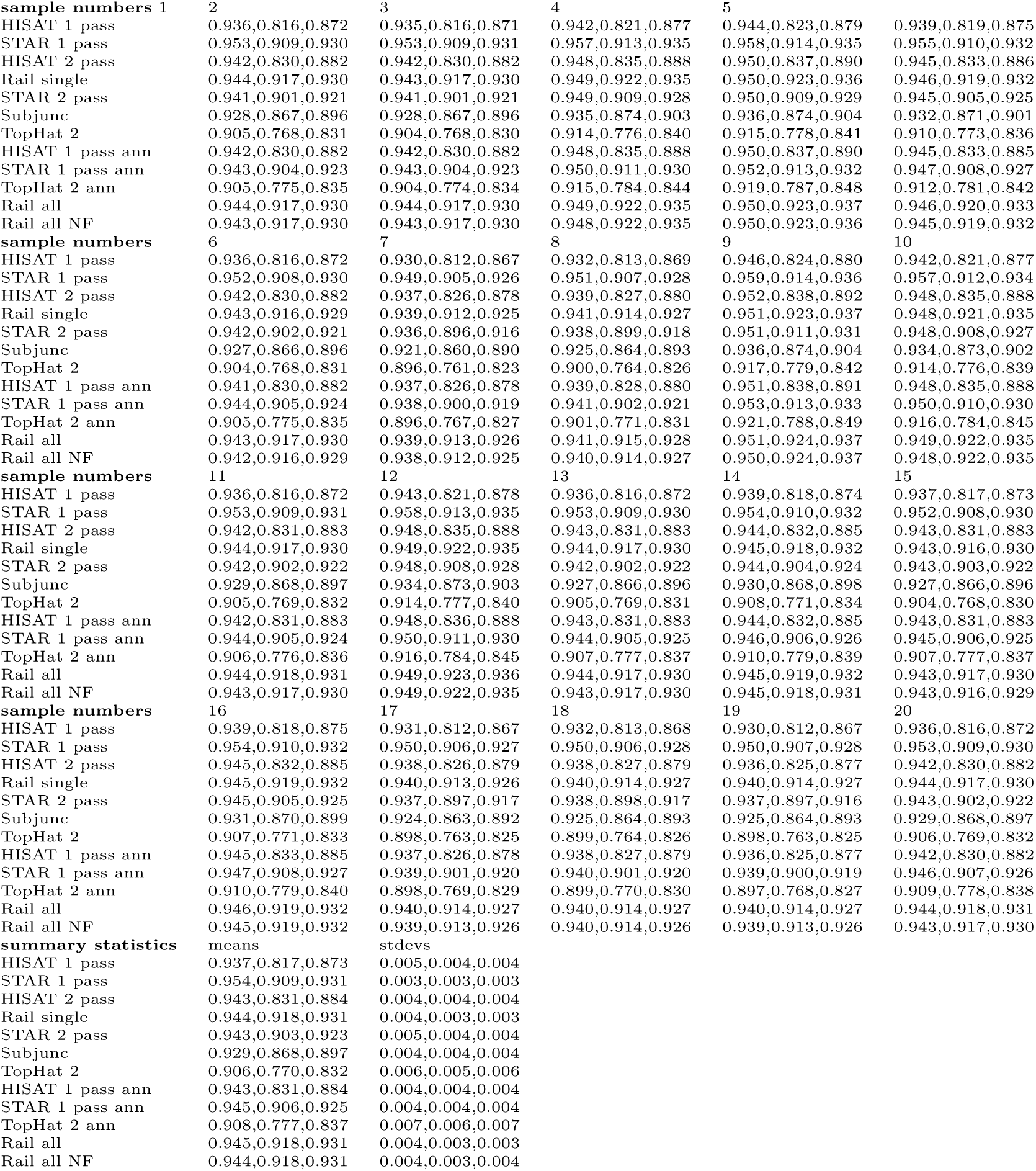
Mapping accuracy for various alignment protocols described in Sections 2.3 and 5.10. Each table item is a tuple in the format “precision, recall, F-score”; these quantities are defined in Section 5.11. Sample numbers correspond to 20 randomly selected samples of the 112 samples we simulated, and they also appear in Table 2. Summary statistics are computed across the 20 samples. Note that the “Rail all” and “Rail all NF” protocols were each run on all 112 simulated samples to augment exon-exon junction calling.

| sample numbers 1 | 2 | 3 | 4 | 5 |  |
| --- | --- | --- | --- | --- | --- |
| HISAT 1 pass | 0.936,0.816,0.872 | 0.935,0.816,0.871 | 0.942,0.821,0.877 | 0.944,0.823,0.879 | 0.939,0.819,0.875 |
| STAR 1 pass | 0.953,0.909,0.930 | 0.953,0.909,0.931 | 0.957,0.913,0.935 | 0.958,0.914,0.935 | 0.955,0.910,0.932 |
| HISAT 2 pass | 0.942,0.830,0.882 | 0.942,0.830,0.882 | 0.948,0.835,0.888 | 0.950,0.837,0.890 | 0.945,0.833,0.886 |
| Rail single | 0.944,0.917,0.930 | 0.943,0.917,0.930 | 0.949,0.922,0.935 | 0.950,0.923,0.936 | 0.946,0.919,0.932 |
| STAR 2 pass | 0.941,0.901,0.921 | 0.941,0.901,0.921 | 0.949,0.909,0.928 | 0.950,0.909,0.929 | 0.945,0.905,0.925 |
| Subjunc | 0.928,0.867,0.896 | 0.928,0.867,0.896 | 0.935,0.874,0.903 | 0.936,0.874,0.904 | 0.932,0.871,0.901 |
| TopHat 2 | 0.905,0.768,0.831 | 0.904,0.768,0.830 | 0.914,0.776,0.840 | 0.915,0.778,0.841 | 0.910,0.773,0.836 |
| HISAT 1 pass ann | 0.942,0.830,0.882 | 0.942,0.830,0.882 | 0.948,0.835,0.888 | 0.950,0.837,0.890 | 0.945,0.833,0.885 |
| STAR 1 pass ann | 0.943,0.904,0.923 | 0.943,0.904,0.923 | 0.950,0.911,0.930 | 0.952,0.913,0.932 | 0.947,0.908,0.927 |
| TopHat 2 ann | 0.905,0.775,0.835 | 0.904,0.774,0.834 | 0.915,0.784,0.844 | 0.919,0.787,0.848 | 0.912,0.781,0.842 |
| Rail all | 0.944,0.917,0.930 | 0.944,0.917,0.930 | 0.949,0.922,0.935 | 0.950,0.923,0.937 | 0.946,0.920,0.933 |
| Rail all NF | 0.943,0.917,0.930 | 0.943,0.917,0.930 | 0.948,0.922,0.935 | 0.950,0.923,0.936 | 0.945,0.919,0.932 |
| sample numbers | 6 | 7 | 8 | 9 | 10 |
| HISAT 1 pass | 0.936,0.816,0.872 | 0.930,0.812,0.867 | 0.932,0.813,0.869 | 0.946,0.824,0.880 | 0.942,0.821,0.877 |
| STAR 1 pass | 0.952,0.908,0.930 | 0.949,0.905,0.926 | 0.951,0.907,0.928 | 0.959,0.914,0.936 | 0.957,0.912,0.934 |
| HISAT 2 pass | 0.942,0.830,0.882 | 0.937,0.826,0.878 | 0.939,0.827,0.880 | 0.952,0.838,0.892 | 0.948,0.835,0.888 |
| Rail single | 0.943,0.916,0.929 | 0.939,0.912,0.925 | 0.941,0.914,0.927 | 0.951,0.923,0.937 | 0.948,0.921,0.935 |
| STAR 2 pass | 0.942,0.902,0.921 | 0.936,0.896,0.916 | 0.938,0.899,0.918 | 0.951,0.911,0.931 | 0.948,0.908,0.927 |
| Subjunc | 0.927,0.866,0.896 | 0.921,0.860,0.890 | 0.925,0.864,0.893 | 0.936,0.874,0.904 | 0.934,0.873,0.902 |
| TopHat 2 | 0.904,0.768,0.831 | 0.896,0.761,0.823 | 0.900,0.764,0.826 | 0.917,0.779,0.842 | 0.914,0.776,0.839 |
| HISAT 1 pass ann | 0.941,0.830,0.882 | 0.937,0.826,0.878 | 0.939,0.828,0.880 | 0.951,0.838,0.891 | 0.948,0.835,0.888 |
| STAR 1 pass ann | 0.944,0.905,0.924 | 0.938,0.900,0.919 | 0.941,0.902,0.921 | 0.953,0.913,0.933 | 0.950,0.910,0.930 |
| TopHat 2 ann | 0.905,0.775,0.835 | 0.896,0.767,0.827 | 0.901,0.771,0.831 | 0.921,0.788,0.849 | 0.916,0.784,0.845 |
| Rail all | 0.943,0.917,0.930 | 0.939,0.913,0.926 | 0.941,0.915,0.928 | 0.951,0.924,0.937 | 0.949,0.922,0.935 |
| Rail all NF | 0.942,0.916,0.929 | 0.938,0.912,0.925 | 0.940,0.914,0.927 | 0.950,0.924,0.937 | 0.948,0.922,0.935 |
| sample numbers | 11 | 12 | 13 | 14 | 15 |
| HISAT 1 pass | 0.936,0.816,0.872 | 0.943,0.821,0.878 | 0.936,0.816,0.872 | 0.939,0.818,0.874 | 0.937,0.817,0.873 |
| STAR 1 pass | 0.953,0.909,0.931 | 0.958,0.913,0.935 | 0.953,0.909,0.930 | 0.954,0.910,0.932 | 0.952,0.908,0.930 |
| HISAT 2 pass | 0.942,0.831,0.883 | 0.948,0.835,0.888 | 0.943,0.831,0.883 | 0.944,0.832,0.885 | 0.943,0.831,0.883 |
| Rail single | 0.944,0.917,0.930 | 0.949,0.922,0.935 | 0.944,0.917,0.930 | 0.945,0.918,0.932 | 0.943,0.916,0.930 |
| STAR 2 pass | 0.942,0.902,0.922 | 0.948,0.908,0.928 | 0.942,0.902,0.922 | 0.944,0.904,0.924 | 0.943,0.903,0.922 |
| Subjunc | 0.929,0.868,0.897 | 0.934,0.873,0.903 | 0.927,0.866,0.896 | 0.930,0.868,0.898 | 0.927,0.866,0.896 |
| TopHat 2 | 0.905,0.769,0.832 | 0.914,0.777,0.840 | 0.905,0.769,0.831 | 0.908,0.771,0.834 | 0.904,0.768,0.830 |
| HISAT 1 pass ann | 0.942,0.831,0.883 | 0.948,0.836,0.888 | 0.943,0.831,0.883 | 0.944,0.832,0.885 | 0.943,0.831,0.883 |
| STAR 1 pass ann | 0.944,0.905,0.924 | 0.950,0.911,0.930 | 0.944,0.905,0.925 | 0.946,0.906,0.926 | 0.945,0.906,0.925 |
| TopHat 2 ann | 0.906,0.776,0.836 | 0.916,0.784,0.845 | 0.907,0.777,0.837 | 0.910,0.779,0.839 | 0.907,0.777,0.837 |
| Rail all | 0.944,0.918,0.931 | 0.949,0.923,0.936 | 0.944,0.917,0.930 | 0.945,0.919,0.932 | 0.943,0.917,0.930 |
| Rail all NF | 0.943,0.917,0.930 | 0.949,0.922,0.935 | 0.943,0.917,0.930 | 0.945,0.918,0.931 | 0.943,0.916,0.929 |
| sample numbers | 16 | 17 | 18 | 19 | 20 |
| HISAT 1 pass | 0.939,0.818,0.875 | 0.931,0.812,0.867 | 0.932,0.813,0.868 | 0.930,0.812,0.867 | 0.936,0.816,0.872 |
| STAR 1 pass | 0.954,0.910,0.932 | 0.950,0.906,0.927 | 0.950,0.906,0.928 | 0.950,0.907,0.928 | 0.953,0.909,0.930 |
| HISAT 2 pass | 0.945,0.832,0.885 | 0.938,0.826,0.879 | 0.938,0.827,0.879 | 0.936,0.825,0.877 | 0.942,0.830,0.882 |
| Rail single | 0.945,0.919,0.932 | 0.940,0.913,0.926 | 0.940,0.914,0.927 | 0.940,0.914,0.927 | 0.944,0.917,0.930 |
| STAR 2 pass | 0.945,0.905,0.925 | 0.937,0.897,0.917 | 0.938,0.898,0.917 | 0.937,0.897,0.916 | 0.943,0.902,0.922 |
| Subjunc | 0.931,0.870,0.899 | 0.924,0.863,0.892 | 0.925,0.864,0.893 | 0.925,0.864,0.893 | 0.929,0.868,0.897 |
| TopHat 2 | 0.907,0.771,0.833 | 0.898,0.763,0.825 | 0.899,0.764,0.826 | 0.898,0.763,0.825 | 0.906,0.769,0.832 |
| HISAT 1 pass ann | 0.945,0.833,0.885 | 0.937,0.826,0.878 | 0.938,0.827,0.879 | 0.936,0.825,0.877 | 0.942,0.830,0.882 |
| STAR 1 pass ann | 0.947,0.908,0.927 | 0.939,0.901,0.920 | 0.940,0.901,0.920 | 0.939,0.900,0.919 | 0.946,0.907,0.926 |
| TopHat 2 ann | 0.910,0.779,0.840 | 0.898,0.769,0.829 | 0.899,0.770,0.830 | 0.897,0.768,0.827 | 0.909,0.778,0.838 |
| Rail all | 0.946,0.919,0.932 | 0.940,0.914,0.927 | 0.940,0.914,0.927 | 0.940,0.914,0.927 | 0.944,0.918,0.931 |
| Rail all NF | 0.945,0.919,0.932 | 0.939,0.913,0.926 | 0.940,0.914,0.926 | 0.939,0.913,0.926 | 0.943,0.917,0.930 |
| summary statistics | means | stdevs |  |  |  |
| HISAT 1 pass | 0.937,0.817,0.873 | 0.005,0.004,0.004 |  |  |  |
| STAR 1 pass | 0.954,0.909,0.931 | 0.003,0.003,0.003 |  |  |  |
| HISAT 2 pass | 0.943,0.831,0.884 | 0.004,0.004,0.004 |  |  |  |
| Rail single | 0.944,0.918,0.931 | 0.004,0.003,0.003 |  |  |  |
| STAR 2 pass | 0.943,0.903,0.923 | 0.005,0.004,0.004 |  |  |  |
| Subjunc | 0.929,0.868,0.897 | 0.004,0.004,0.004 |  |  |  |
| TopHat 2 | 0.906,0.770,0.832 | 0.006,0.005,0.006 |  |  |  |
| HISAT 1 pass ann | 0.943,0.831,0.884 | 0.004,0.004,0.004 |  |  |  |
| STAR 1 pass ann | 0.945,0.906,0.925 | 0.004,0.004,0.004 |  |  |  |
| TopHat 2 ann | 0.908,0.777,0.837 | 0.007,0.006,0.007 |  |  |  |
| Rail all | 0.945,0.918,0.931 | 0.004,0.003,0.003 |  |  |  |
| Rail all NF | 0.944,0.918,0.931 | 0.004,0.003,0.004 |  |  |  |

### 5.13 Exon-exon junction coverage bias

**Figure 7:**
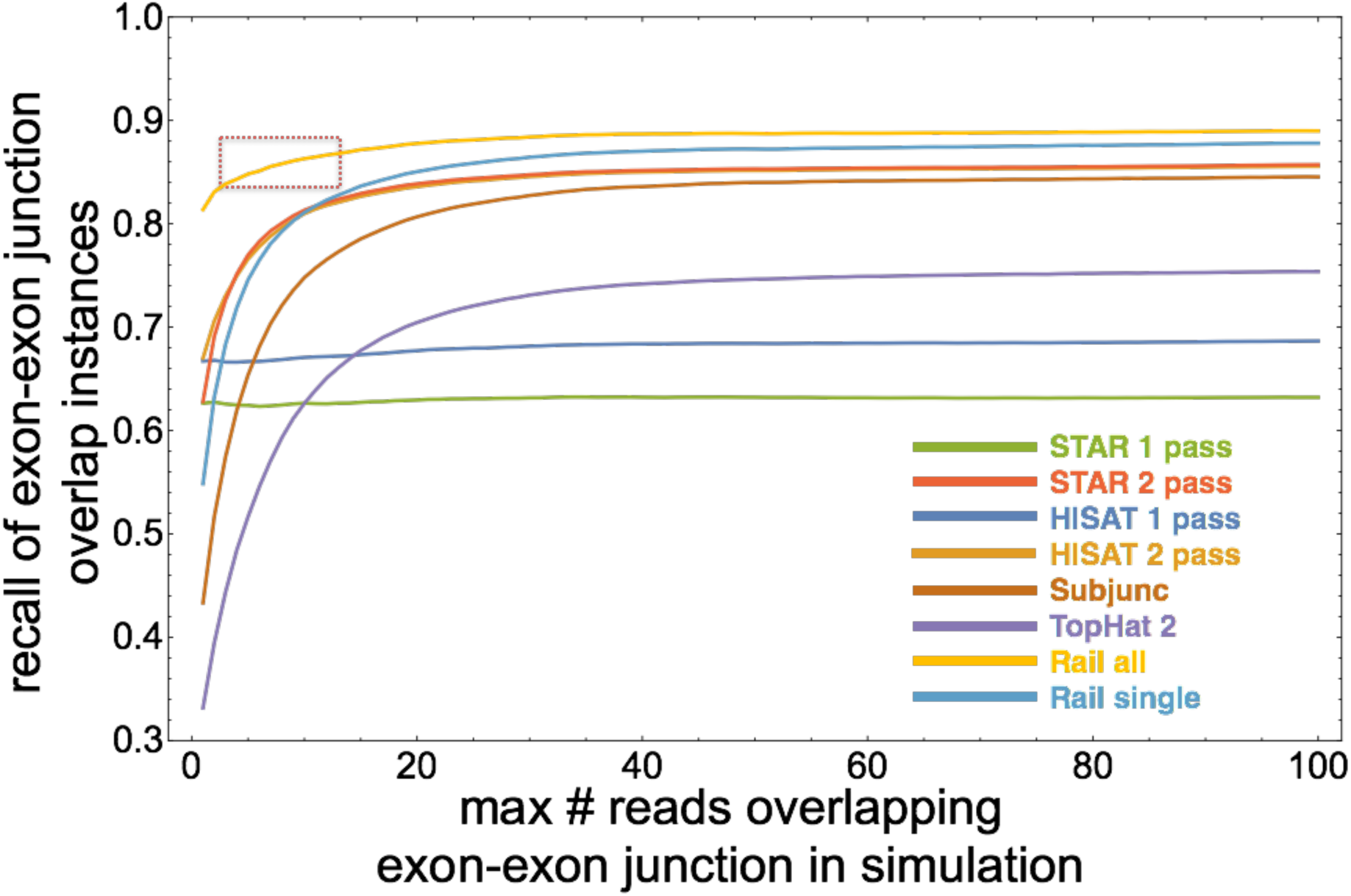
Recall of exon-exon junction overlap instances for unannotated protocols run on the simulated sample “NA20768 female TSI HMGU 5-1-1.” The “Rail all” protocol (yellow line at the top) achieves high recall without succumbing significantly to coverage bias where exon-exon junctions are covered by few reads (red dotted box), unlike 2-pass single-sample protocols “STAR 2 pass,” “HISAT 2 pass,” Subjunc, TopHat 2, and “Rail single.” 1-pass protocols are not subject to this exon-exon junction coverage bias but do not achieve high recall.

Let *c*(*r*) denote the true coverage of the minimally covered exon-exon junction overlapped by a read **r** in a sample *s*. Here, “true coverage” refers to the number of reads truly derived from loci overlapping a given exon-exon junction. For example, if **r** overlaps two exon-exon junctions *i*_1_ and *i*_2_, and *i*_1_ is truly covered by 10 reads in *S* from *s* while *i*_2_ is truly covered by 25 reads in *S*, *c*(*r*) = 10. Figure 7 plots recall for various protocols that do not use annotation run on a randomly selected simulated sample (“NA20768_female_TSI_HMGU_5-1-1”) at various thresholds *t* of *c*(*r*); a given point on any curve accounts for every read **r** for which *c*(*r*) ≤ *t*. This plot reveals a bias present in certain spliced alignment protocols that involve realignment (i.e., the 2 pass protocols): the correct alignments of reads that overlap exon-exon junctions whose true coverage is low are recalled less often. This exon-exon junction coverage bias has a simple explanation: if there are more reads covering a given exon-exon junction, it is likelier that exon-exon junction is detected initially (on the first pass) by an alignment protocol. So during realignment, it is likelier that splice junctions are detected successfully for reads overlapping highly covered exon-exon junctions than it is for poorly covered exon-exon junctions. By contrast, if no realignment is performed, recall should be independent of *c*(*r*): the probability a splice junction is detected for a given read is independent of the probabilities splice junctions are detected in other reads. Neither “STAR 1 pass” nor “HISAT 1 pass” perform realignment, so their recalls are more or less horizontal lines across the plot. However, recall is diminished compared to protocols that do perform realignment. (HISAT achieves higher recall than STAR on one pass because it accumulates an exon-exon junction list as it aligns a sample, searching for a larger number of exon-exon junctions as more reads are aligned. But the aligner does not have a large list of exon-exon junctions at its disposal when it is working through the first reads, so one pass still cannot match the recall of two passes.) Figure 8 plots recall for various protocols that use annotation as well as the “Rail all” protocol at various thresholds *t* of *c*(*r*). “STAR 1-pass ann,” “HISAT 1-pass ann,” and “TopHat 2 ann” do indeed provide high recall while minimizing exon-exon junction coverage bias; an annotation can reveal the locations of poorly covered exon-exon junctions. *However, annotation-based protocols are themselves subject to a bias*: it is likelier that exon-exon junctions listed in the annotation are found than it is that novel exon-exon junctions are. This bias is not discernible in Figure 8. Moreover, because the annotated protocols tested here are given the artificial advantage of foreknowledge of the entire space of exon-exon junctions sampled by the simulated samples, Figure 8 exaggerates the positive effect of annotation. “Rail all” achieves significant exon-exon junction coverage bias mitigation, however, faring nearly as well as even the annotated protocols: note that recall on the y-axis never drops below 0.79.

**Figure 8:**
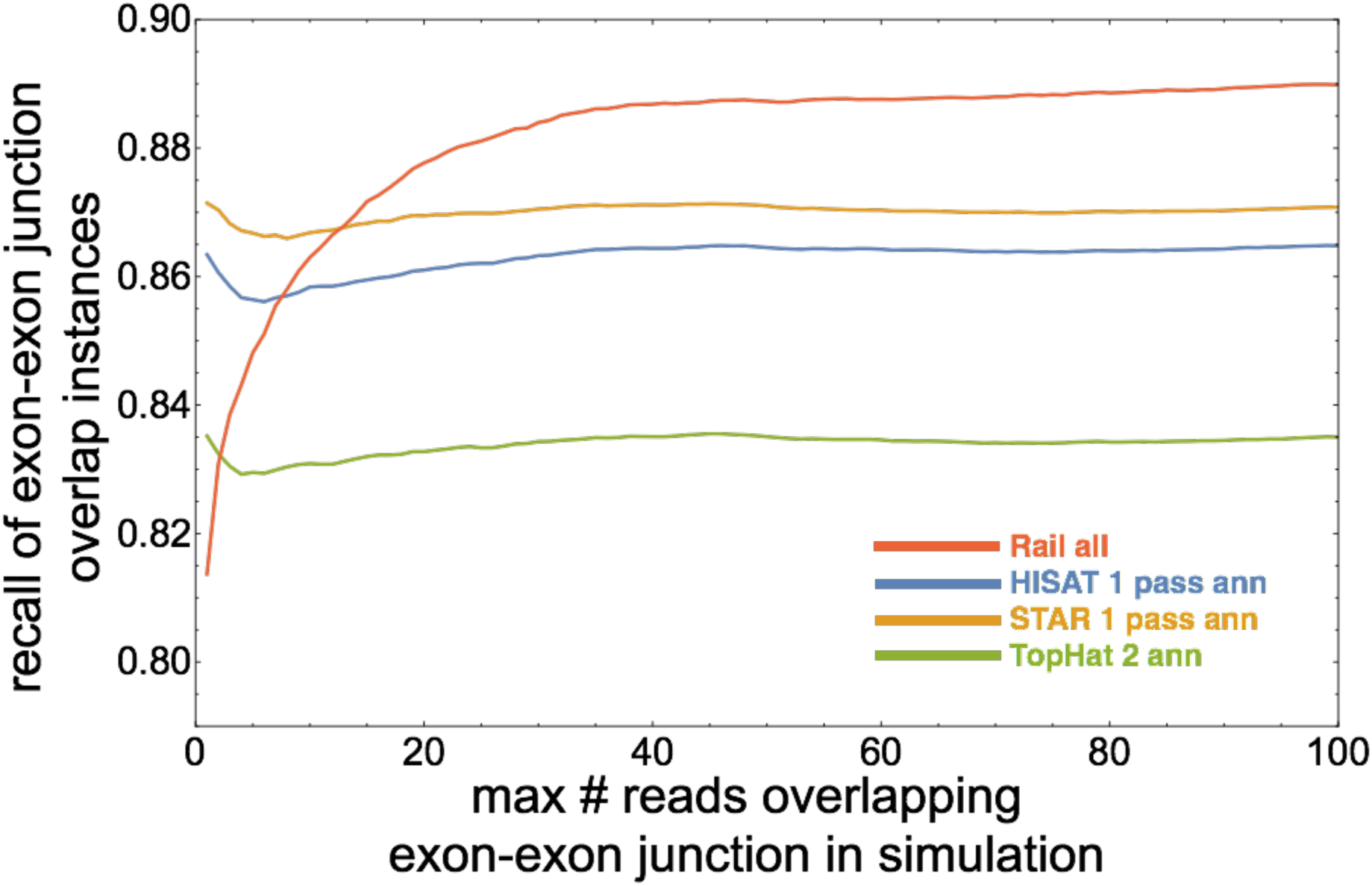
Recall of exon-exon junction overlap instances annotated protocols run on the simulated sample “NA20768 female TSI HMGU 5-1-1.” The “Rail all” protocol (red curved line) does not appear to correct for exon-exon junction coverage bias as well as annotated protocols, but annotated protocols have an artificial advantage: they are given as additional input every true exon-exon junction sampled in simulation.

**The “Rail all” protocol corrects for annotation bias while minimizing exon-exon junction coverage bias and simultaneously providing high recall through realignment.** The protocol benefits from how there is a higher probability any given exon-exon junction—even one that is poorly covered across samples—is detected. When many samples are analyzed together, the exon-exon junction coverage bias is greatly diminished.

### 5.14 Wall-clock times of alignment protocols on a single computer and a single sample

We measured wall-clock times of the various alignment protocols studied in the main text on a single computer using 8 and 16 2.4-GHz Intel Xeon E5-2665 processors. We invoked the Linux commandline utility `taskset` to confine processor use to exactly 8 or 16 cores and ran each alignment protocol on the randomly selected paired-end GEUVADIS sample whose SRA accession number is ERR205018, which has 44,703,518 75-bp reads (counting ends of paired-end reads separately). The “STAR 1 pass ann” and “TopHat 2 ann” protocols require construction of indexes from gene annotation. We used the Gencode v12 annotation [40] and timed index construction separately. Rail-RNA can suppress some outputs, saving end-to-end computation time. We measured Rail-RNA’s performance writing both all its default outputs (alignment BAMs, coverage bigWigs, indel and junction BEDs, and cross-sample TSVs) as well as writing only alignment BAMs. Our results are in Table 6. Note that the operating configurations tested here are not the preferred way to run Rail-RNA, which is designed to take advantage of hundreds of processing cores in a distributed environment and eliminate redundant alignment work across many samples.

**Table 6:** Wall-clock times of various alignment protocols for GEUVADIS sample ERR205018 run on 8 and 16 2.4-GHz Intel Xeon E5-2665 processing cores. The annotated protocols “STAR 1 pass ann” and “TopHat 2 ann” require onetime index construction from gene annotation, and we report wall-clock times with and without index construction using the Gencode v12 annotation. Rail-RNA’s wall-clock time was measured with default outputs and with only BAM outputs.

|  | 8 cores | 16 cores |
| --- | --- | --- |
| HISAT 1 pass ann | 4m26s | 3m1s |
| HISAT 1 pass | 4m33s | 3m13s |
| HISAT 2 pass | 8m57s | 6m20s |
| Rail-RNA (BAM only) | 103m55s | 59m25s |
| Rail-RNA (all default outputs) | 118m46s | 64m40s |
| STAR 1 pass ann (with index time) | 46m48s | 37m22s |
| STAR 1 pass ann (no index time) | 5m38s | 3m59s |
| STAR 1 pass | 4m6s | 2m24s |
| STAR 2 pass | 10m14s | 7m35s |
| Subjunc | 19m54s | 14m44s |
| TopHat 2 ann (with index time) | 239m34s | 224m54s |
| Topat 2 ann (no index time) | 132m54s | 98m53s |
| TopHat 2 | 110m38s | 84m27s |

### 5.15 Expressed Regions analysis

For each sample, base level coverage is imported from the bigWig files and then adjusted by the total number of mapped reads to represent the coverage from a library size of 80 million reads. This adjustment could have been left out, and we performed it only to make base-level coverage take intuitively manageable values. Expressed regions (ERs) are then determined by the `regionMatrix()`function from `derfinder` (version 1.0.10) [43] applied to the adjusted coverage with a mean cutoff of 5 reads. ERs are identified for each chromosome separately and then merged with a custom script. Overlap with Ensembl v75 [40] known exons, exon-exon junctions, and intergenic regions is determined with the corresponding genomic state object via `makeGenomicState()` and `annotateRegions()` from `derfinder`.

`regionMatrix()` also generates the corresponding adjusted coverage matrix for library size of 80 million reads assuming a read length of 75 base pairs. This matrix is log_2_ transformed using an offset of 1 and is available via Figshare [54]. A linear regression is then fitted with the 15 technical variables available: population, RIN value, RNA extraction batch, RNA concentration, RNA quantity used, library preparation date, primer index, method concentration measure, library concentration, library size, library concentration used, cluster kit, sequencing kit, cluster density, and sequencing lane. The percent of variance explained is determined by the residual sum of squares for the given variable divided by the total sum of squares from all variables as well as the residual variation.

The GEUVADIS expressed regions are available in `supplementaryExpressedRegions.csv.` This CSV file has the following columns:

1. seqnames: chromosome where the ER is located.
2. start: extreme left position of the ER.
3. end: extreme right position of the ER.
4. width: width of the ER in base pairs.
5. strand: strand of the ER.
6. value: mean adjusted coverage for the ER across all samples.
7. area: sum of adjusted coverage for the ER across all samples.
8. indexStart: internal index start position.
9. indexEnd: internal index end position.
10. cluster: cluster ID for the ER where ERs belong to the same cluster if they are at most 3000 base pairs apart. IDs are unique by chromosome only.
11. clusterL: cluster length in base pairs.

### 5.16 Rail-RNA’s objective function

Rail-RNA seeks the best alignment for each input read as follows. Let [*n*] denote {1,…,n} for some positive integer *n*. Say Rail-RNA finds a number *N* of possible alignments of a given read **r**. Let *i∈* [*N*] index these alignments. Each has an alignment score *s*(*i*) and a number *n*(*i*) of exon-exon junctions overlapped by the alignment. *s*(*i*) is determined by Bowtie 2’s local alignment parameters [33], which assign penalties to gaps and mismatches in a user-configurable fashion. Consider the set of reads in the same sample as **r** such that each has exactly one highest-scoring alignment. Call these the uniquely aligned reads. Let *c*(*i, j*) be the number of uniquely aligned reads covering the *j*th junction overlapped by alignment *i*, and suppose

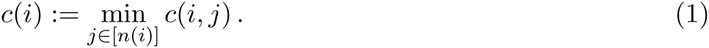

If *n*(*i*) = 0, set *c*(*i*) = 1. Let

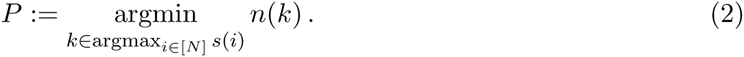

*P* will often have one element. When there is more than one, Rail-RNA draws the primary alignment of **r** at random weighted by *c*(*i*):

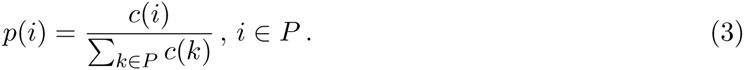

### 5.17 Implementation details

Rail-RNA is built on the MapReduce programming model [30], which uses an economy of fundamental abstractions to promote scalable cluster computing. A problem is divided into a sequence of computation and aggregation steps. Each step consumes a set of key-value pairs and outputs a set of key-value pairs. A step is itself divided into a collection of embarrassingly parallel tasks to be executed concurrently by workers under the aegis of a scheduler. An aggregation step places pairs with the same key in the same partition and sorts each partition by value. A computation step is either a reduce step or a map step; a reduce step involves prior aggregation, while a map step does not. Each worker may be assigned several tasks. A worker in a reduce step operates on a sorted partition at a time, where a given task may be comprised of several such partitions. A worker in a map step, on the other hand, operates on key-value pairs in no particular order. Each map or reduce step consumes a set of key-value pairs and outputs a set of unordered key-value pairs.

All three of Rail-RNA’s modes—elastic, parallel, and single-computer—run the same set of Python scripts using PyPy [55], a fast Python interpreter. In parallel and single-computer modes:

- A master Python script runs as an independent process and operates as a task scheduler; that is, it assigns tasks to worker processes on the same computer (in single-computer mode) or on other computers (in parallel mode) and ensures that all of a step’s tasks are complete before moving on to the next step.
- A computation step is performed by having each worker stream a (possibly sorted) group of input files into a PyPy process running a Python script.
- In general, the result of either a map step or a reduce step is a set of output files, each containing key-value pairs and each the result of a different worker. If the next step is a map step, each worker streams a different output file into a PyPy process running the step’s Python script. If the next step is a reduce step, the output files require prior aggregation. This is achieved as follows. Each of the previous step’s output files is streamed into a different worker, which hashes on each key of a given key-value pair in the stream to assign the pair to a different task (file). Each worker thus outputs several files, one for each task, and there are typically as many tasks as there are workers. Moreover, after all workers are done hashing on keys, there may be many files corresponding to the same task. Each of these files is next assigned to a different worker, which calls GNU Coreutils sort (the standard command-line sort utility that comes with most UNIX-based operating systems) to sort the key-value pairs in the file so that pairs in the same partition are adjacent and sorted by value in the order required by the reduce step. Finally, groups of sorted files corresponding to the same task are assigned to the same worker, which merges them in sorted order and streams the result into a PyPy process running the reduce step’s Python script.

Elastic mode uses Hadoop [29] via Amazon Elastic MapReduce [56] to manage task scheduling and aggregation. We use Hadoop Streaming, a tool that comes with Hadoop, to run our external Python scripts spanning all computation steps using PyPy. The elastic-mode implementation of Rail-RNA is largely managed by Elastic MapReduce: it requires that we specify

- the number and type of EC2 instances to provision for assembling a Hadoop computer cluster.
- a job flow encoding the order of map and reduce steps as well as where on S3 they can find their inputs and write their outputs.

### 5.18 Detail: Preprocess reads

In local and parallel modes, workers divide the reads for each sample (typically stored in one FASTQ or a pair of FASTQs) among several gzip-compressed files. Splitting a sample’s reads up in this manner enables their distribution among workers in the next step of the pipeline. In elastic mode, the reads for each sample are written to a single LZO-compressed file that may subsequently be indexed for the same purpose as file-splitting in local and parallel modes: this also permits fast distribution of file splits among workers in the next step. The distribution is facilitated by Twitter’s Elephant Bird library [57] for Hadoop.

In local and parallel modes, if all input reads are on a locally accessible filesystem, the preprocess map step described in the main text is preceded by two load-balancing steps. In the first—itself a map step—each worker operates on a sample at a time, counting the number of reads. Only one worker is active in a step that follows, collecting read counts across samples and deciding how many and which reads should be processed by a given worker at a time in the preprocess map step. This ensures that load is distributed evenly across workers during preprocessing. In elastic mode or if some samples must be downloaded from the Internet, a worker operates on a sample at a time in the preprocess map step. This permits deletion of a given downloaded set of reads by a worker as soon as it is preprocessed, saving storage space.

Importantly, if a read is from a paired-end sample, its mate’s nucleotide sequence is encoded in the read’s name. This way, if the read is found to have more than one highest-scoring alignment in the next step, the mate sequence can easily be retrieved to help resolve the tie.

### 5.19 Detail: Align reads to genome

Reads are partitioned by nucleotide sequence so that each worker can operate on a set of reads that share the same sequence at once. A task is composed of several such sets; to reduce redundancy in analysis, a worker nominates from each set a read with the highest mean quality score in that set for alignment using Bowtie 2 [33] in its “local” mode. (To avoid redundancy between sequences and reversed complements, a read’s nucleotide sequence is taken to be either the original or its reversed complement, whichever is first in lexicographic order.) Call the set of these nominated reads *A*_1_. If a read in *A*_1_ has exactly one perfect alignment, we assume it does not overlap any exon-exon junctions and place it *and all other reads with the same nucleotide sequence* in set *F*; that is, the alignment found is assigned to all reads that share the same nucleotide sequence. If a read is from a single-end sample and has more than one perfect alignment, we also assume the read does not overlap any exon-exon junctions and place all it in set *F*; however, we place other reads with the same nucleotide sequence in set *A*_2_. If a read in *A*_1_ is from a paired-end sample and has more than one alignment, whether or not the alignment is perfect, we place the read and all other reads with the same nucleotide sequence in set *A*_2_. Reads in *A*_2_ are aligned on a second run of Bowtie 2 performed in-step by the same worker. This lets Bowtie 2 break ties in their alignment scores to select primary alignments, now with paired-end data where available. Alignments of reads in *F* are final and appear in Rail-RNA’s terminal output; they attain the highest possible alignment score and overlap no exon-exon junctions.

Bowtie 2’s local mode soft-clips the ends of alignments if doing so improves local alignment score. We exploit this feature to determine which read sequences should be probed for overlapped exon-exon junctions. If the primary alignment of any read in *A*_1_ has at least one edit or is softclipped, we add the read to a set *C*, place its associated nucleotide sequence in set *S*, and add all other reads with the same nucleotide sequence to *A*_2_. Note that *S* is composed of unique read sequences, while *A*_1_, *A*_2_, and *F* may contain reads with the same sequence. Any read whose nucleotide sequence is in *S* is realigned in a later step. Set *S* is further divided into subset *S*_search_ and its complement *S*_no search_. If the primary alignment of a read in *A*_1_ lacks a soft-clipped end spanning at least some user-specified number of bases (by default 1), its nucleotide sequence is added to *S*_no search_. Otherwise, the sequence is added to *S*_search_. In subsequent steps, read sequences from *S*_search_ are searched for exon-exon junctions and realigned to references containing transcript fragments—contiguous exonic base sequences from the genome. Reads whose nucleotide sequences are in *S*_no search_ are never searched for exon-exon junctions but are also realigned to transcript fragments.

Primary alignments of reads in *A*_2_ are found on a second run of Bowtie 2 in local mode. If a primary alignment is perfect, its associated read is promoted to *F*; if it contains at least one edit, the read is placed in *C*. Alignments of reads in *C* are saved for comparison with realignments of these reads to transcript fragments in later steps.

Every unique read sequence in *S* is hashed to place it one of *P* × *N*_*s*_ “index” bins, where *N*_*s*_ is the number of samples under analysis in the Rail-RNA run and *P* is the number of FM indexes that will be built per sample in a future realignment step. More specifically, all reads whose sequences are in the same bin are aligned to the same index in the step described in Section 3.9. Every unique read sequence in *S* is also “readletized”: it is divided into short overlapping segments, called readlets. In the next step, readlets are aligned to the genome so any introns that appear between alignments of successive nonoverlapping readlets to the same contig can be inferred. **Information about in which samples each read sequence occurs is passed on to subsequent steps.**

**Figure 9:**
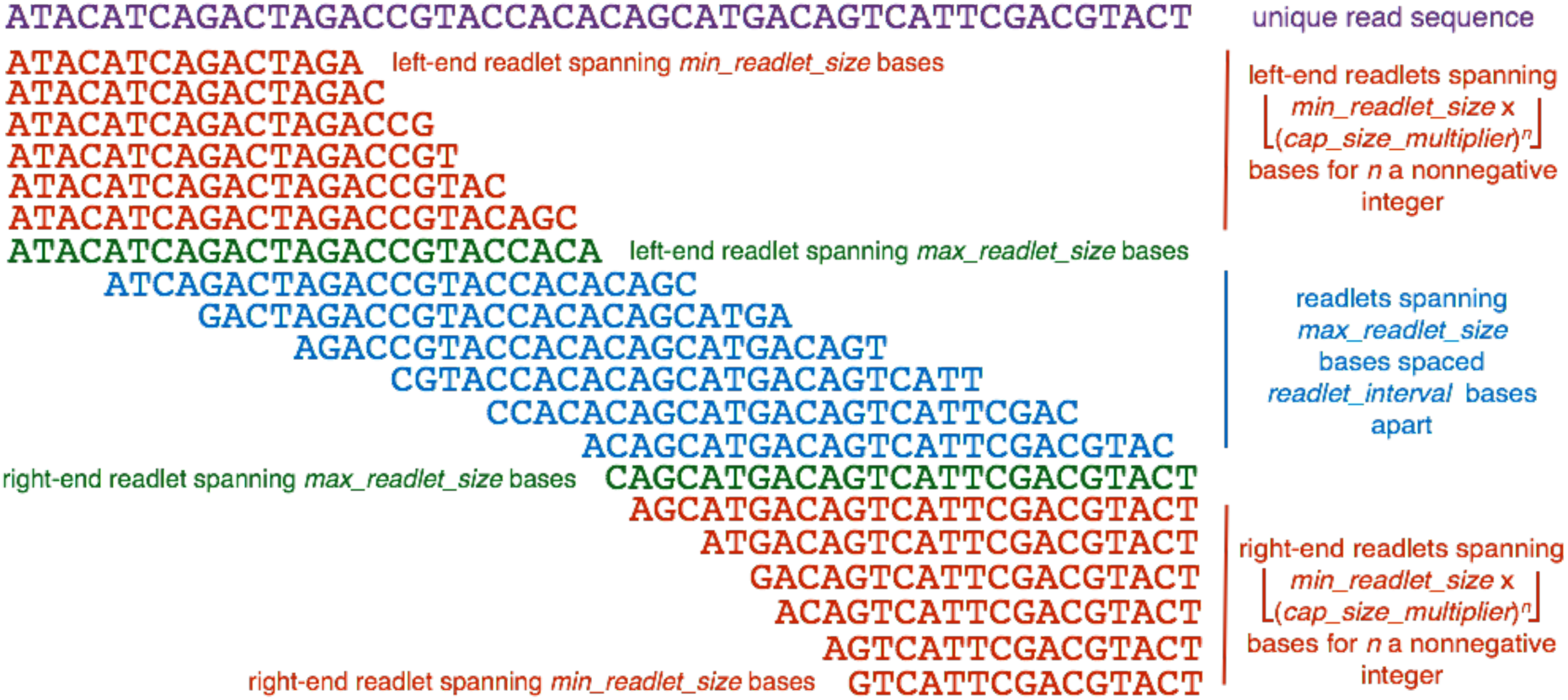
Rail-RNA’s scheme for readletizing (segmenting) a read sequence into short subsequences for alignment to the genome to infer exon-exon junction positions. In this example, *min_readlet_size* is 15, *max_readlet_size* is 25, *cap_size_multiplier* is 1.1, and *readlet_interval* is 4.

The readletizing scheme is detailed in Figure 9. There is always a readlet at each end of the read spanning *max_readlet_size* bases. Intervening readlets spanning *max_readlet_size* bases are spaced *readlet interval* bases apart. There are also smaller readlets (“capping readlets”) at each end of the read. Their sizes are spanned by the set

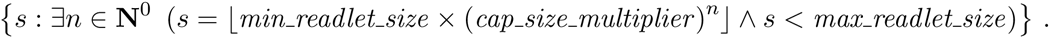

Parameters from the previous paragraph are positive integers constrained as follows.

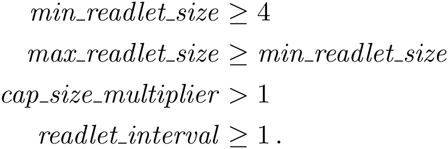

Their default values of, respectively, 15, 25, 1.1, and 4 may be toggled by the user.

### 5.20 Detail: Detect exon-exon junctions using readlet alignments

To explain Rail-RNA’s algorithm for searching for exon-exon junctions, we restrict our attention to a single read **r** and its readlets.^2^ To be precise in our discussion, we use the (slightly redundant) notation (**d**, *p***_*r*_**, *s, p*_*s*_, ℓ, *k*) to denote the *k*th *exact match to the reference* of a readlet **d** spanning *ℓ* bases that is displaced *p*_**r**_ bases away from the left end of **r**; the alignment is to a position displaced *p*_*s*_ bases away from the left end of strand *s* of the reference assembly. When two readlet alignments (**d**_1_, *p*_**r**,1_, *s*_1_, *p*_*s*_1__, _1_, *ℓ*_*1*_ *k*_1_), (**d**_2_, *p*_***r***_, *s*_2_, *p*_*s*_*2*__,_*2*_, *ℓ*_2_, *k*_2_) are *compatible*:

1. **d**_1_ ≠ **d**_2_; the alignments are not of the same readlet.
2. *s*_1_ = *s*_2_ = *s* for some strand *s*; the two readlets align to the same strand.
3. *p*_**r**,1_ *< p*_**r**,2_ *⇔ p*_*s,*1_ *< p*_*s,*2_; if their positions along the read are different, the readlets are ordered in the same way along the read as their alignments along *s*.
4. *p*_**r**,1_ = *p*_**r**,2_ *⇔ p*_*s,*_1 = *p*_*s,*2_; if their positions along the read are the same, the readlets align to the same position along *s*.

**Figure 10:**
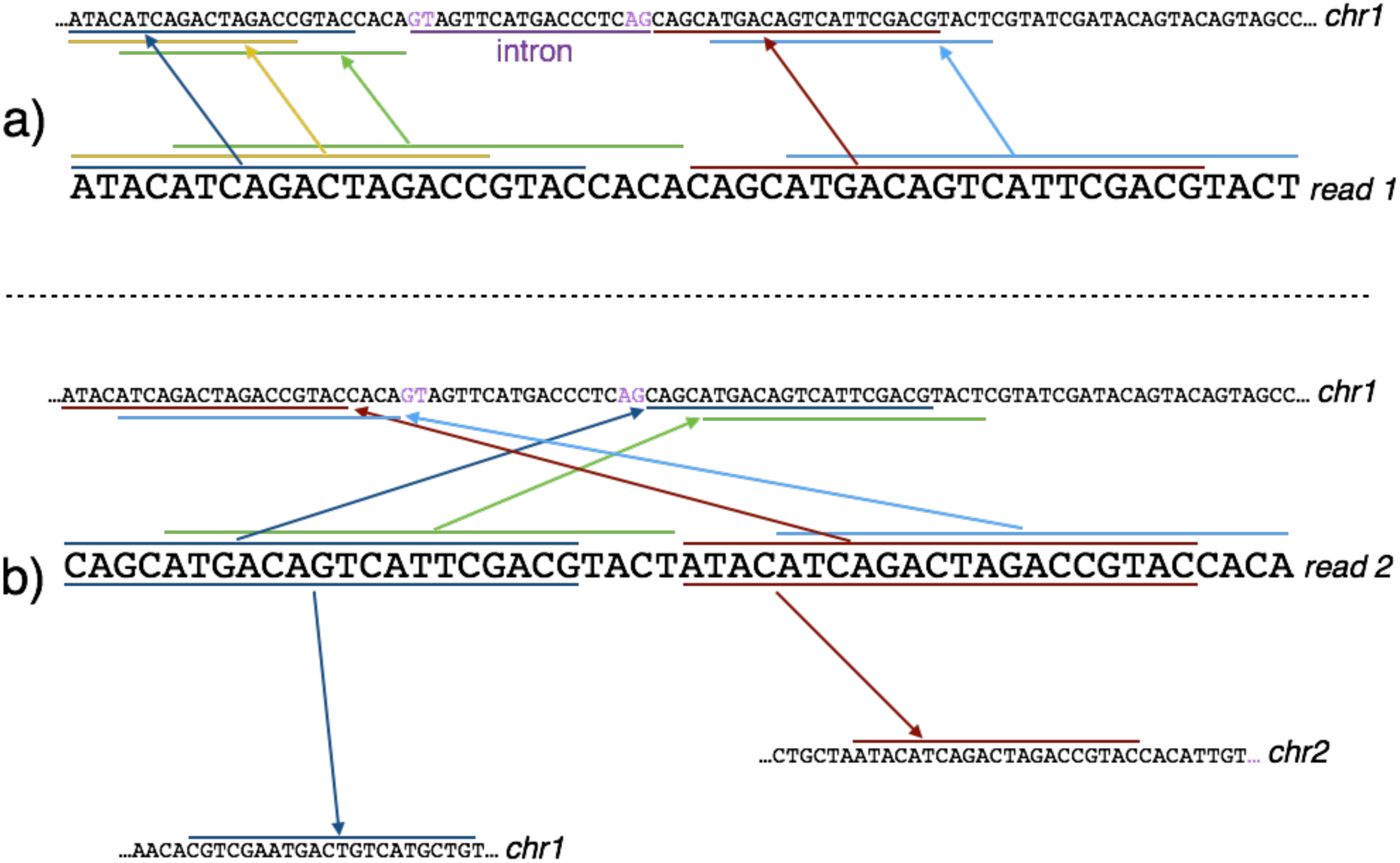
a) Ideally, every readlet of a read aligns uniquely to the same strand of a contig (e.g., the forward strand of chr1). b) Often, the ideal case depicted in a) is not realized and one or more of the following occurs: readlets do not align in the order in which they appear along the read (top); readlets align to multiple strand (the readlets that are underlined and overlined with arrows pointing to their alignments); and readlets align in the right order along the same contig but to different strands. (Note that the readlet mapping at the bottom left of the figure illustrates the reversed complement of a readlet aligning to the forward strand.)

Ideally, the readlet alignments derived from **r** are mutually compatible, as depicted in the top half of Figure 10: all of read 1’s mapped readlets align uniquely to the same strand (here, the forward strand of *chr1*) in the order in which they appear along the read; and further, if two readlets begin at the same position along the read, they align to the same position along the reference. Suppose *i* ∈ [*N*] indexes a set *R* = {(**d**_*i*_, *p*_**r**,*i*_, *s, p*_*s,i*_, *ℓ*_*i*_, *k*_*i*_)} of mutually compatible readlet alignments. Note that each alignment in *R* corresponds to a distinct readlet. Let *S*(*i*) = {*j* : ∃ *j* ∈ [*N*] *p*_**r**,*i*_ = *p*_**r**,*j*_}; that is, *S*(*i*) indexes the set of readlets in *R* whose displacements from the left end of **r** are the same as **d**_*i*_’s. When two readlet alignments (**d**_*m*_, *p*_**r**,*m*_, s, p_*s,*1_, ℓ _m_), (**d**_*q*_, *p*_**r**,*q*_, *s, p*_s,q_, ℓ _2_) *2 R* are *consecutive*:

1. *p*_**r**,*m*_ ≠ *p*_**r**,*q*_; the readlets do not occur at the same position along the read.
2. (*ℓ* _*m*_ *∈* argmax_*j∈S*(*m*)_ *ℓ* _*j*_) *^* (*ℓ* _*q*_ *∈* argmax_*j∈(q)*_ *ℓ* _*j*_) holds; each readlet is the longest mapped readlet occurring at its respective position along the read.
3. (*p*_**r**_,_*m*_ < *p*_**r**,*q*_ ⇒ {*i* : ∃*i ∈* [*N*] *p*_**r**_,*m* < *p*_**r**_,*i* < *p*_**r**_,*q}* = Ø) *^* (*p*_**r**,*q* < p__**r**_,*m* ⇒ *{i* : ∃*i ∈* [*N*] *p*_**r**_,*q < p*_**r**_,*i* < *p*_**r**_,*m*} = Ø) holds; there is no mapped readlet between **d**_*m*_ and **d**_*q*_ exclusive.

An example of two consecutive readlet alignments is provided by the subsequences overlined and underlined in green and brown in the top half of Figure 10. These alignments also illustrate one way an exon-exon junction overlapped by a read is recovered by Rail-RNA: when two readlets align consecutively to the reference on either side of exactly one intron, the presence of an exon-exon junction is inferred by searching for characteristic two-base motifs denoting its donor and acceptor sites. In order of descending prevalence in mammalian genomes according to a survey of GenBank annotated genes [58], the (donor, acceptor) site pairs Rail-RNA recognizes are (GT, AG), (GC, AG), and (AT, AC). Such sites are reproduced in the reference sequence if the sense strand is the forward strand; the (GT, AG) combination appears in Figure 10. If the sense strand is the reverse strand, the (acceptor, donor) signals may still be read off the reference from left to right as (CT, AC), (CT, GC), and (GT, AT).

**Figure 11:**
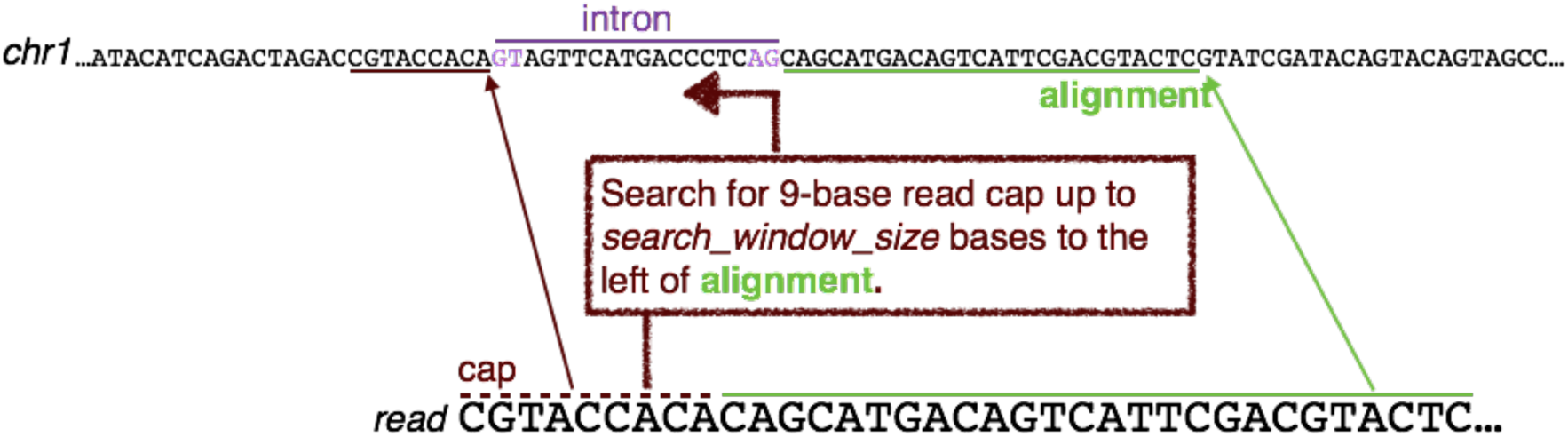
Rail-RNA searches for a match to the read cap that overlaps no aligned readlets if the cap spans at least *min exon size* bases, where *min exon size* is by default 9.

Figure 11 displays a case where a read has an unmapped 9-base “cap” to the left of a readlet aligned by Bowtie in *R*: the nine bases do not belong to any mapped readlet. In this event, Rail-RNA searches the reference *search window size* bases upstream of the leftmost readlet alignment for maximum matching prefixes of the cap. By default, *search window size* is 1000. If a prefix found spans fewer than *min exon size* bases, it is ignored. by default, *min exon size* is 9. Otherwise, Rail-RNA treats the maximum matching prefix closest to the leftmost readlet alignment as a readlet alignment itself. This procedure is mirrored if there is a cap at the right end of the read, where the search is then downstream of the rightmost aligning readlet. So the set of mutually compatible readlet alignments *R* may be augmented by up to two caps.

Assuming a pair of consecutive readlet alignments is correct and that only one intron lies between them, they uniquely determine the length *L* of the intron along the reference in the absence of intervening indels whose net length is nonzero: for consecutive readlets

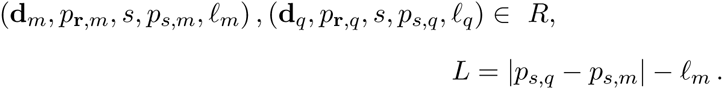

If more than one intron lies between two consecutive readlet alignments, *L* as defined above is the *sum* of the lengths of the introns. For every pair of consecutive readlet alignments in *R*, Rail-RNA searches for up to two intervening introns with valid (donor, acceptor) motifs that satisfy the constraint on *L*. When searching for two introns, Rail-RNA requires that the size of the exon between them is above an adjustable parameter *min exon size*. Denote as *F* the region of the reference framed by the two consecutive readlets. Search windows are confined to the *search window size* bases at either end of *F*. Several candidate introns or intron pairs may be uncovered within *F* and in the vicinity of its ends. Each candidate is placed in one of three preference classes according to (donor, acceptor) signal: (GT, AG) is preferred to (GC, AG), which is preferred to (AT, AC). Rail-RNA also ranks each candidate within its respective class: its score is determined from global alignment of the read subsequence overlapped by the two consecutive readlet alignments to the exonic bases surrounding the candidate intron or intron pair. Here, we assign +1 for each matched base and −1 for each single-base gap and mismatched base. Rail-RNA selects candidate introns and intron pairs with the highest score in the most-preferred class with at least one alignment. For every intron, a (key, value) pair is written for each sample in which **r** occurs. More specifically, the output of this step is:

**key:** strand of origin of intron, intron start position, intron end position

**value:** a list of tuples [(*i,; β*_*i*_)], where *i* indexes a sample and; *β*_*i*_ is the number of reads in the sample with the searched read sequence in which the intron was found.

The discussion in this section is so far predicated on how all the readlet alignments derived from a read **r** are mutually compatible, forming a valid set *R*. This ideal situation is not always realized. The bottom half of Figure 10 displays how readlet alignments may be incompatible: readlets may not align to the same strand in the same order in which they appear along the read, or a readlet may have more than one alignment. In such cases, Rail-RNA selects a “consensus” set of mutually compatible alignments from all the readlet alignments as follows. Consider the set *V* = *{*(**d**_*i*_, *p***_*r*_**,*i, s, p*_*s,i*_, *ℓ*_*i*_, *k*_*i*_)} of all alignments of all readlets reported by Bowtie. Form a complete signed graph *𝒢* = (*V, E*) whose vertices correspond to readlet alignments:

1. Start with the vertices disconnected.
2. Place a “+” edge between every pair of alignments that are the closest compatible alignments of their corresponding readlets. Here, “closest” means “having the smallest number of bases between their start positions along the reference.”
3. Complete the graph with “*-*” edges.

Correlation clustering finds the clustering of vertices that maximizes the number of agreements— that is, the sum of the number of “+” edges within clusters and the number of “*-*” edges between clusters. A consensus set of compatible alignments may be derived from the largest such cluster. However, correlation clustering is an NP-hard problem [59], so we use a randomized 3-approximation algorithm introduced in [60]. Its pseudocode is given in Algorithm 1. After this procedure, the pivot alignment in each cluster *C*_*i*_ is compatible with every other alignment in that cluster. However, it is not guaranteed that all the alignments in a given *C*_*i*_ are mutually compatible. So Rail-RNA sorts the clusters in order of descending size and loops through it: it first forms a unweighted graph *𝒢*_*i*_ for a given *C*_*i*_, placing an edge between two alignments if and only if they are compatible. An efficient variant of the Bron-Kerbosch algorithm [61] is then used to obtain a maximum clique *C*_*i*_ for *C*_*i*_. This algorithm enumerates all *maximal* cliques, which includes all *maximum* cliques. If the algorithm finds more than one maximum clique for a given cluster, the maximum clique is chosen at random. A good explanation of the algorithm is provided in the following direct quote from [62].

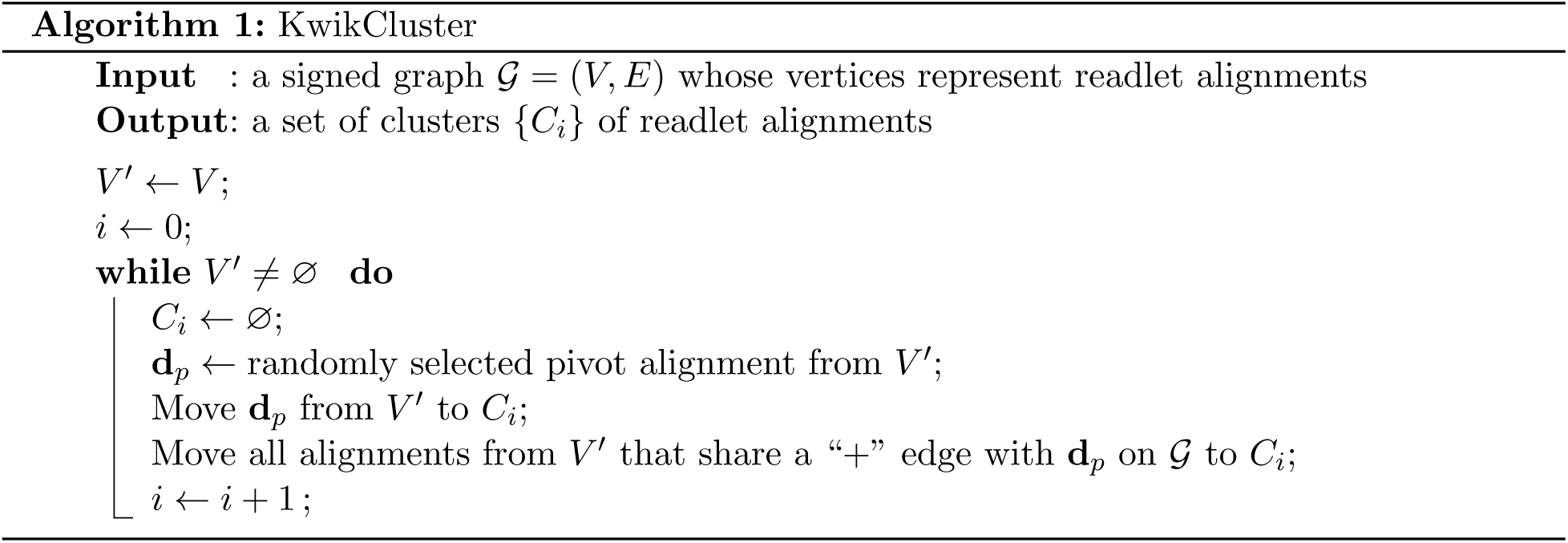

> A recursive call to the Bron–Kerbosch algorithm provides three disjoint sets of vertices *R*, *P*, and *X* as arguments, where *R* is a (possibly non-maximal) clique and *P ⋃ X* = r(*R*) are the vertices that are adjacent to every vertex in *R*. The vertices in *P* will be considered to be added to clique *R*, while those in *X* must be excluded from the clique; thus, within the recursive call, the algorithm lists all cliques in *P ⋃ R* that are maximal within the subgraph induced by *P ⋃ R ⋃ X*. The algorithm chooses a candidate *v* in *P* to add to the clique *R*, and makes a recursive call in which *v* has been moved from *R* to *P*; in this recursive call, it restricts *X* to the neighbors of *v*, since non-neighbors cannot affect the maximality of the resulting cliques. When the recursive call returns, *v* is moved to *X* to eliminate redundant work by further calls to the algorithm. When the recursion reaches a level at which *P* and *X* are empty, *R* is a maximal clique and is reported…. To list all maximal cliques in the graph, this recursive algorithm is called with *P* equal to the set of all vertices in the graph and with *R* and *X* empty.

If *𝓒*_*i*_ spans more vertices than *C*_*i*+1_—the next cluster from the sorted list—or if *C*_*i*_ is the last cluster from the list, *and if there are no cliques of the same size among those computed so far*, Rail-RNA selects *𝓒*_*i*_ as the set of mutually compatible readlet alignments from which it infers exon-exon junctions. If there are multiple cliques of the same size, the software does not search the read sequence for exon-exon junctions. So when there are no such ties, Rail-RNA selects the largest clique from among the {*C*_*i*_}. While the time complexity of the Bron-Kerbosch algorithm is exponential in the number of graph vertices, it executes quickly in practice: 1) it is run only on each cluster of readlet alignments where at least one alignment is compatible with other alignments from the cluster, so there are few vertices; and 2) the number of possible of alignments Rail-RNA studies for a given readlet is limited by Bowtie’s `-m` parameter; as mentioned above, no readlet alignments are reported when more than by default 30 alignments are uncovered.

### 5.21 Spliced alignment using substrings and seed-and-extend

Rail-RNA’s strategy of performing spliced alignment by first extracting substrings from reads and searching for where they occur in the reference genome using an index, is very well studied. It was used in early spliced alignment tools including QPALMA [16] and TopHat [9]. Rail-RNA’s strategy, including its us of overlapping substrings, is perhaps most similar to the seed-and-vote strategy of Subjunc [38]. In seed-and-vote, equally spaced, overlapping 16-bp segments of reads called subreads (synonymous with readlets) are mapped to the reference genome with a hash index, which requires no mismatched bases between reference and read sequence. Each alignment of a subread is counted as a vote for its location in the reference. A single subread may thus vote for up to as many locations in the genome as it has alignments. If the two mapping locations with the most votes are on the same strand, their intervening sequence is then searched for the (donor, receptor) signal (GT, AG) to identify an exon-exon junction. Some salient differences between Rail-RNA’s algorithm and Subjunc’s seed-and-vote follow.

- Rail-RNA’s algorithm never defines boundaries of mapping locations of groups of readlet alignments when resolving the alignments of multimapping readlets. Rather, it obtains clusters of mutually compatible readlet alignments, where compatibility is illustrated in Figure 10. The largest cluster of mutually compatible readlet alignments is nominated for a search for exon-exon junctions.
- Seed-and-vote searches for (donor, acceptor) signals between the top two mapping locations, while the largest cluster of readlet alignments found by Rail-RNA may include groups of alignments to the same strand in more than two disconnected regions. Such a cluster may thus capture more than two distinct mapping locations, each contained in a different exon, and a read may be found to overlap two or more exon-exon junctions on first pass.
- Rail-RNA searches for (donor, receptor) signals (GT, AG), (GC, AG), and (AT, AC), while Subjunc searches for (GT, AG) only (when it is not also searching for fusion events). Searching for more (donor, receptor) signals increases recall at the expense of precision.

The crux of the difference between Rail-RNA’s strategy and seed-and-vote is that the largest cluster of readlet alignments found by Rail-RNA—the “nominated” cluster—may include several mapping locations whose number is not specified *a priori*, while Subjunc searches explicitly for the top two mapping locations.

### 5.22 Stringency of the junction filter

As described in the main text, Rail-RNA applies a junction filter to remove poorly supported exon-exon junctions from consideration prior to realignment. By default, a junction survives the filter if it is (a) covered by 5 or more reads in one sample, or (b) covered by at least 1 read in 5% of samples. So a read covered by exactly 1 read in 6 out of 100 samples will survive the filter, but a read covered by exactly 4 reads in 4 out of 100 samples (and not covered in any other sample) will be removed.

The filter’s behavior depends on the exon-exon junction profile across samples, but it is instructive to consider it in relation to simpler filters. We use the name “coverage-1” to describe a filter that keeps any junction covered by at least one read in any sample. We use “coverage-5” to describe a filter that keeps any junction covered by at least 5 reads in any sample. The “coverage-5” filter is more stringent than the “coverage-1” filter.

The Rail-RNA filter, which we call “borrow-5-5,” has a stringency that is intermediate between “coverage-1” and “coverage-5.” In the case where no junction is covered in 5% of samples, the “borrow-5-5” filter is the same as the “coverage-5 filter.” In the case where every junction covered in one sample is also covered in at least 5% of samples, the “borrow-5-5” filter is the same as the “coverage-1” filter.

Real RNA-seq datasets are between these extremes. If the samples are replicates of the same condition, we expect gene expression profiles and junction profiles to be fairly consistent across samples, similarly to the scenario where “borrow-5-5” acts like the less stringent “coverage-1” filter. If the samples are very different, then “borrow-5-5” acts like the more stringent “coverage-5” filter.

### 5.23 Detail: Enumerate intron configurations along read segments

Intron configurations are obtained as follows. A directed acyclic graph (DAG) is constructed for every combination of sample and strand for which introns were detected. Each vertex of the DAG represents a unique intron *k*_*i*_ = (*s*_*i*_, *e*_*i*_), where *s*_*i*_ is the coordinate of the first base of the intron, and *e*_*i*_ is the coordinate of the first base that follows the intron. An edge occurs between two introns *k*_1_ and *k*_2_ if and only if they do not overlap—that is, the coordinates spanned by the introns are mutually exclusive—and no intron *k*_3_ occurs between *k*_1_ and *k*_2_ such that *k*_1_, *k*_2_, and *k*_3_ do not overlap. An edge always extends from an intron with smaller coordinates to an intron with larger coordinates. Each edge is weighted by the number of exonic bases between the introns it connects. Figure 12 depicts a portion of an example DAG.

**Figure 12:**
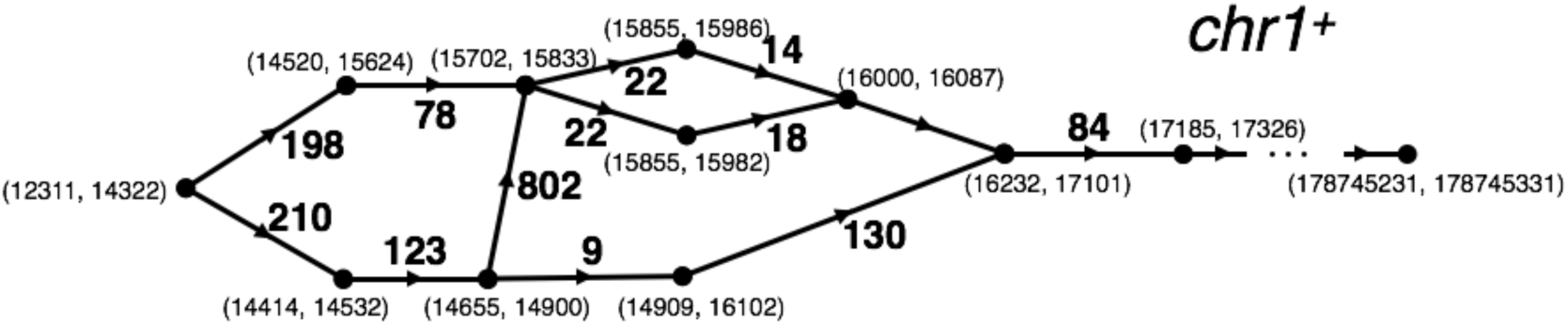
An example DAG. Vertices correspond to introns and are labeled by their start and end positions. An edge extends from one intron to another that comes after it along the strand. Every edge is weighted by the number of exonic bases between the introns it connects.

The paths through the DAG span all possible combinations of nonoverlapping introns along the strand for a given sample. Finding all subpaths (sequences of introns), each of whose weights is less than *readlet config size*, enumerates all possible combinations of exon-exon junctions a read segment **s**(*readlet_config_size*) can overlap. In fact, the parameter *readlet_config_size* could theoretically be set to equal the maximum length of all reads across samples. Then, after concatenating and indexing the exonic sequences surrounding intron configurations to form transcript fragments (isofrags) and indexing, Bowtie 2 could realign entire reads to isofrags end-to-end to immediately obtain candidate spliced alignments. Unfortunately, finding intron combinations in “hot spots,” where there are many alternative splicings and short exons, can become computationally intractable for large *readlet_config_size*. Even taking *readlet_config_size* = 50 can become challenging in later steps, substantially increasing the time it takes to build a Bowtie 2 index of transcript fragments. This explains our choice *readlet_config_size* = 35: it’s large enough so Bowtie 2 can successfully identify which read sequences are derived from which transcript fragments, soft-clipping as necessary, but also small enough to make intron configuration enumeration and subsequent index construction manageable.

A worker handles one given strand for a given sample at a time. Enumerating combinations of exon-exon junctions that could be overlapped by read segments proceeds for each (sample, strand) combination. Rail-RNA uses a streaming algorithm that alternates between **construction** and **consumption** of the DAG. Only a portion of the DAG is in memory at a time. Call this portion the mDAG. For a given worker, the input to the construction-consumption algorithm is a sorted list of introns detected by Rail-RNA in a given sample along a given strand, where the sort key is intron start position. The construction operation reads a chunk of this list and augments the mDAG. A subsequent consumption operation erases a part of the mDAG after the intron combinations for that part have been enumerated. Construction and consumption operations are described in detail below.

**Construction:** The edges of the DAG are output for consumption in an order consistent with a topological ordering of its vertices. In other words, an edge is not generated until every edge for which its source vertex is a sink vertex has already been generated. As mentioned above, the construction subroutine operates on a streamed list of intron positions [{(*s*_*i*_, *e*_*i*_)}] sorted in order of ascending *s*_*i*_, as provided by a previous aggregation step. Two data structures are needed to encode the DAG as it is constructed: the set *U*, containing intron vertices that do not yet have any children, and *L*, a dictionary that, where possible, maps each intron (the key) to its corresponding successive nonoverlapping intron *with the smallest end position read so far* (the value). **For each** new intron *k*_*i*_ = (*s*_*i*_, *e*_*i*_) read:

1. The vertices in *U* and *L* are checked for whether *k*_*i*_ is their child—that is, whether vertices previously read are connected to *k*_*i*_—and corresponding edges are **yield**ed. As edges are yielded, vertices from *U* may be promoted to *L*.
2. Each “value” vertex *k*_*j*_ in *L* is replaced with *k*_*i*_ if *e*_*i*_ *< e*_*j*_.
3. Every vertex *k*_*j*_ in *L* is checked for whether its corresponding value is itself a key. If this condition is satisfied, an edge can *never* connect *k*_*j*_ with any introns streamed later, and *k*_*j*_ is removed from *L*.

Rail-RNA generates the DAG 10, 000, 000 bases at a time. After a portion of the DAG is produced, all possible exon-exon junction combinations that can be overlapped by a read segment **s**(*readlet_config_size*) are **output for processing in the next steps.** Rail-RNA accomplishes this by considering each vertex separately and finding the ways its associated exon-exon junction can be the first along **s**; that is, it walks every path starting from that vertex edge by edge until its weight exceeds *readlet_config_size*.

**Consumption:** The mDAG is dynamic: edges and vertices may have previously been consumed so that there are new source vertices and sink vertices. A source vertex *k* is removed from the mDAG when all paths originating at every child vertex *k*_*i*_ of *k* have been reviewed to find intron configurations that correspond to exon-exon junctions spanned by *readlet_config_size* exonic bases. *k* can be removed at this time because it is no longer needed to find by how many exonic bases to the left of *k*_*i*_ an isofrag can possibly extend before running into an intron. More specifically, an edge from *k* to *k*_*i*_ is removed if and only if:

1. All edges with *k*_*i*_ as the source have been generated. By construction, this occurs if *k*_*i*_ has at least one grandchild.
2. Every path of maximal length originating at *k*_*i*_ = (*s*_*i*_, *e*_*i*_) whose weight is less than (*readlet_config_size* 1) can be constructed. Suppose *{*(*s*_*j*_, *e*_*j*_)*}* are the children of *k*_*i*_, and let *{s, e}* be the as-yet-unconsumed vertex with the largest *s* from the dynamic DAG. This occurs if for every j, *s e_j_ > readlet_config_size*−1−(*s*_*i*_−*e*_*j*_).

The alternation continues until the end of the strand is reached. At this point, the edges connecting remaining vertices in *U* and their new sinks are **yield**ed, and **remaining intron configurations are output.**

### 5.24 Detail: Retrieve and index transcript fragments that overlap exon-exon junctions

Consider the key-value pair ((*t, {*(*s*_*i*_, *e*_*i*_)*}*), (*l*_*j*_, *r*_*j*_)), where *i ∈* [*N*] and *j* indexes samples. The key is an intron configuration composed of *N* introns, where the variable *t* stores the strand of the intron. The value is an ordered pair (*l*_*j*_, *r*_*j*_). The variable *l*_*j*_ stores the number of bases that must be traversed upstream of *s*_1_ before another complete intron is found along *s* in some sample; the variable *r*_*j*_ stores the number of bases that must be traversed downstream of *e*_*N*_ before another complete intron is found along *s* in some sample. If no intron precedes (*s*_1_, *e*_1_), *l* is recorded as “not available,” and if no intron succeeds (*s*_*N*_, *e*_*N*_), *r* is recorded as “not available.”

In the first step referenced in the main text, each worker operates on an intron configuration at a time. In general, an intron configuration will have appeared in several samples, but the *{*(*l*_*j*_, *r*_*j*_)*}* could vary from sample to sample because the introns identified in each sample are different. During the step, a given transcript fragment is assembled from reference exonic bases surrounding the intron configuration (*s, {*(*s*_*i*_, *e*_*i*_)*}*); the fragment starts at coordinate *s*_1_ min_*j*_ *l*_*j*_ and terminates at coordinate *e*_1_ + min_*j*_ *r*_*j*_. Because we use the minima of *l*_*j*_ and *r*_*j*_, there is no chance a given transcript fragment overlaps exons besides those demarcated by the introns (*t, {*(*s*_*i*_, *e*_*i*_)*}*). Each transcript fragment is associated with a new reference name encoding its start and end coordinates and which junctions are overlapped.

### 5.25 Detail: Finalize combinations of exon-exon junctions overlapped by read sequences

Local alignment of a given read sequence to isofrags with Bowtie 2 in general outputs alignments of several (possibly overlapping) read segments. The positions of the alignments along the isofrags gives precise information about where and which exon-exon junctions could possibly be overlapped by the read sequence. An undirected graph is constructed from this list of exon-exon junctions, each of which corresponds to a distinct intron. Each vertex is a different intron (*t, s, e*), where *t* denotes strand, *s* the start coordinate on the reference, and *e* the end coordinate. An edge is placed between two vertices (*t*_*i*_, *s*_*i*_, *e*_*i*_) and (*t*_*j*_, *s*_*j*_, *e*_*j*_) if and only if

1. *t*_*i*_ = *t*_*j*_; the introns are on the same strand.
2. There are no more than *L* 1 exonic bases between introns *i* and *j*.
3. min(*e*_*i*_, *e*_*j*_)–max(*s*_*i*_, *s*_*j*_) *≥* 0; the introns do not overlap.

The Bron-Kerbosch algorithm is executed on this graph to determine all its maximal cliques. This algorithm is described toward the end of Section 5.20. Maximal cliques are considered candidate combinations of exon-exon junctions that may be overlapped by the read sequence.

### 5.26 Detail: Compile coverage vectors and write bigWigs

We describe how we write bigWigs encoding coverage of the genome by the reads in each sample here. Previous steps that have decided primary alignments (Sections 3.2 and 3.10) have output *exon differentials* associated with each read. The exon differentials of a read are assigned according to the following rules.

1. Divide each contig of a reference genome up into partitions, each spanning *genome partition length* bases. Rail-RNA takes *genome partition length* = 5000 by default.
2. Associate a +1 with the start coordinate of every exonic chunk of the read’s primary alignment. Here, exonic chunks are separated by exon-exon junctions or, optionally, deletions from the reference.
3. Associate a − 1 with the end position of every exonic chunk of the read’s primary alignment.
4. Associate a +1 with the beginning of every genome partition spanned by the read.

These rules permit the reconstruction of the coverage of every base in a given genome partition *solely from the exon differentials within that partition*. This is conceptually simpler and requires less computational effort than compiling coverage from *intervals* representing the spliced alignments.

Figure 13 illustrates how coverage can be reconstructed from differentials. Initialize a variable *v* recording coverage at a given base to 0, and walk across a given partition base by base from left to right. Add the exon differentials at a given base to *v* to obtain the coverage at that base. Note that one exonic chunk lies across the border between partitions 2 and 3 and thus contributes an extra exon differential +1 to the beginning of partition 3. This special case illustrates the necessity of Rule 4: the extra +1 is necessary to obtain the correct coverage value of 1 at the beginning of the partition.

In the initial reduce step *R*_1_, each worker operates on all of the exon differentials for a given sample and genome partition. The worker computes that sample’s coverage at each base in the partition, a task made easier because the differentials are pre-sorted along the length of the partition. The output is compacted using run-length encoding: coverage is only written at a given position if it has changed from the previous position.

A challenge in this step is load balance. The number of exon differentials to be tallied can vary drastically across genome partitions. Therefore, an additional reduce step *R*_1*a*_ precedes *R*_1_. *R*_1*a*_ simply sums exon differentials at a given position in the genome for a given sample. The way key/value pairs are paritioned in preparation for *R*_1*a*_, a given task will process a random subset of genome positions. So for *R*_1*a*_, the distribution of load across workers is close to uniform in practice. With the differentials pre-summed in this way, the number of key-value pairs processed in a given partition in step *R*_1_ is upper-bounded by the number of positions in the partition, greatly improving load balance in that step.

**Figure 13:**
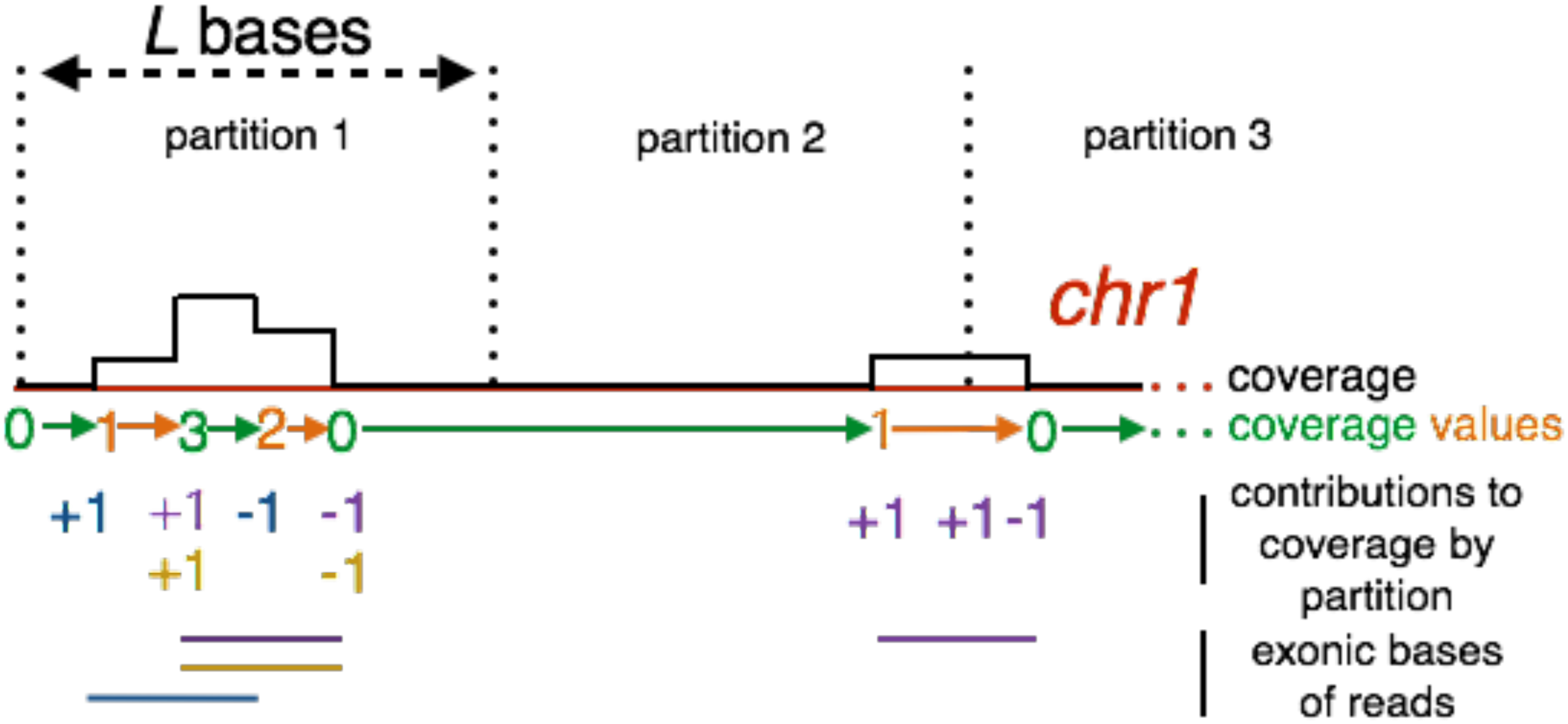
An illustration of how exon differentials are summed along a partition to give appropriate coverage values. Note that there is an extra +1 at the beginning of partition 3 so a worker operating on only partition 3 can obtain the right initial coverage value.

An extra complexity of our discussion above is that some reads have “unique” primary alignments— that is, each of these reads has exactly one highest-scoring alignment—while other reads have several highest-scoring alignments, and the primary alignment is chosen by breaking the tie. We classify exon differentials according to whether they are derived from uniquely aligning reads and in *R*_1*a*_, obtain the exon differential sum at a given base position for each of primarily alignments and uniquely aligning reads.

The coverage output of *R*_1_ is partitioned by sample and sorted by genome coordinate. In another reduce step *R*_2_, each worker can thus operate on all the coverage output of a given sample at once. Coverage for primary alignments and uniquely aligning reads are each written to a file in bedGraph format [63], which is converted to a compressed bigWig using the command-line tool bedGraphToBigWig [41]. In elastic mode, the bigWigs for each sample are uploaded to S3, where it is subsequently accessible on the Internet. During the step *R*_2_, Rail-RNA also computes the upper-quartile normalization factor [64] associated with each sample.

### 5.27 Analyzing in batches

In some settings, not all samples comprising the dataset of interest are available at the same time. They might become available in “batches” due to study design or to the timing of consortium data releases. In these cases, the user would like to be able to analyze each batch separately without the final results being dependent on the order in which the samples were analyzed. This is sometimes called the *N* + 1 problem.

Rail-RNA allows the user to analyze a set of related samples in batches such that the output depends only on which batch is analyzed first, and not on any other aspect of batch order. By default, Rail-RNA outputs the “isofrag” index for a given run—that is, a Bowtie 2 index of transcript fragments spanning exon-exon junctions. These are inferred from all the samples given to Rail-RNA (see Section 3.7). For a batch of *N* samples where *N* is large, and given filter parameters *K* and *J* (see Sections 3.5 and 5.22), junction content will change only slightly when one additional sample is added.

Rail-RNA permits reuse of the isofrag index from the first run to align subsequent batches. In a Rail-RNA run that reuses the isofrag index, Rail-RNA does not search for exon-exon junctions. This mode of running Rail-RNA is analogous to modes of running alternative alignment software that use a gene annotation, where only junctions from the annotation rather than novel junctions are found. We are currently exploring what appropriate values of *N* are for obtaining stable exon-exon junction indexes for reuse.

## Footnotes

1 Hereafter, we use the symbol $ to denote USD.

2 Here, a read is really a unique read sequence. In this discussion, we take the “left end of a read” to mean the left end of either the sequence or its reversed complement, whichever is first in lexicographic order.

## References

[1]. T. C. Glenn, “Field guide to next-generation dna sequencers,” Molecular Ecology Resources, vol. 11, no. 5, pp. 759–769, 2011.

[2]. E. C. Hayden, “Is the $1,000 genome for real?,” Nature News, 2014.

[3]. Z. Wang, M. Gerstein, and M. Snyder, “Rna-seq: a revolutionary tool for transcriptomics,” Nature Reviews Genetics, vol. 10, no. 1, pp. 57–63, 2009.

[4]. F. Ozsolak and P. M. Milos, “Rna sequencing: advances, challenges and opportunities,” Nature Reviews Genetics, vol. 12, no. 2, pp. 87–98, 2010.

[5]. R. Leinonen, H. Sugawara, and M. Shumway, “The sequence read archive,” Nucleic acids research, p. gkq1019, 2010.

[6]. J. Lonsdale, J. Thomas, M. Salvatore, R. Phillips, E. Lo, S. Shad, R. Hasz, G. Walters, F. Garcia, N. Young, et al., “The genotype-tissue expression (gtex) project,” Nature genetics, vol. 45, no. 6, pp. 580–585, 2013.

[7]. “The Cancer Genome Atlas.” http://cancergenome.nih.gov, Aug. 2013.

[8]. K. Wang, D. Singh, Z. Zeng, S. J. Coleman, Y. Huang, G. L. Savich, X. He, P. Mieczkowski, S. A. Grimm, C. M. Perou, et al., “Mapsplice: accurate mapping of rna-seq reads for splice junction discovery,” Nucleic acids research, p. gkq622, 2010.

[9]. C. Trapnell, L. Pachter, and S. L. Salzberg, “Tophat: discovering splice junctions with rnaseq,” Bioinformatics, vol. 25, no. 9, pp. 1105–1111, 2009.

[10]. D. Kim, G. Pertea, C. Trapnell, H. Pimentel, R. Kelley, and S. L. Salzberg, “Tophat2: accurate alignment of transcriptomes in the presence of insertions, deletions and gene fusions,” Genome Biol, vol. 14, no. 4, p. R36, 2013.

[11]. A. Dobin, C. A. Davis, F. Schlesinger, J. Drenkow, C. Zaleski, S. Jha, P. Batut, M. Chaisson, and T. R. Gingeras, “Star: ultrafast universal rna-seq aligner,” Bioinformatics, vol. 29, no. 1, pp. 15–21, 2013.

[12]. T. Bonfert, G. Csaba, R. Zimmer, and C. C. Friedel, “A context-based approach to identify the most likely mapping for rna-seq experiments,” BMC bioinformatics, vol. 13, no. Suppl 6, p. S9, 2012.

[13]. N. Philippe, M. Salson, T. Commes, and E. Rivals, “Crac: an integrated approach to the analysis of rna-seq reads,” Genome biology, vol. 14, no. 3, p. R30, 2013.

[14]. J. Hu, H. Ge, M. Newman, and K. Liu, “Osa: a fast and accurate alignment tool for rna-seq,” Bioinformatics, vol. 28, no. 14, pp. 1933–1934, 2012.

[15]. Y. Zhang, E.-W. Lameijer, P. AC’t Hoen, Z. Ning, P. E. Slagboom, and K. Ye, “Passion: a pattern growth algorithm-based pipeline for splice junction detection in paired-end rna-seq data,” Bioinformatics, vol. 28, no. 4, pp. 479–486, 2012.

[16]. F. De Bona, S. Ossowski, K. Schneeberger, and G. Rätsch, “Optimal spliced alignments of short sequence reads,” BMC Bioinformatics, vol. 9, no. Suppl 10, p. 07, 2008.

[17]. N. Cloonan, Q. Xu, G. J. Faulkner, D. F. Taylor, D. T. Tang, G. Kolle, and S. M. Grimmond, “Rna-mate: a recursive mapping strategy for high-throughput rna-sequencing data,” Bioinformatics, vol. 25, no. 19, pp. 2615–2616, 2009.

[18]. G. R. Grant, M. H. Farkas, A. D. Pizarro, N. F. Lahens, J. Schug, B. P. Brunk, C. J. Stoeckert, J. B. Hogenesch, and E. A. Pierce, “Comparative analysis of rna-seq alignment algorithms and the rna-seq unified mapper (rum),” Bioinformatics, vol. 27, no. 18, pp. 2518–2528, 2011.

[19]. S. Huang, J. Zhang, R. Li, W. Zhang, Z. He, T.-W. Lam, Z. Peng, and S.-M. Yiu, “Soapsplice: genome-wide ab initio detection of splice junctions from rna-seq data,” Frontiers in genetics, vol. 2, 2011.

[20]. K. F. Au, H. Jiang, L. Lin, Y. Xing, and W. H. Wong, “Detection of splice junctions from paired-end rna-seq data by splicemap,” Nucleic acids research, vol. 38, no. 14, pp. 4570–4578, 2010.

[21]. D. W. Bryant, R. Shen, H. D. Priest, W.-K. Wong, and T. C. Mockler, “Supersplatspliced rna-seq alignment,” Bioinformatics, vol. 26, no. 12, pp. 1500–1505, 2010.

[22]. T. D. Wu and S. Nacu, “Fast and snp-tolerant detection of complex variants and splicing in short reads,” Bioinformatics, vol. 26, no. 7, pp. 873–881, 2010.

[23]. S. Marco-Sola, M. Sammeth, R. Guigo, and P. Ribeca, “The gem mapper: fast, accurate and versatile alignment by filtration,” Nature methods, vol. 9, no. 12, pp. 1185–1188, 2012.

[24]. G. Jean, A. Kahles, V. T. Sreedharan, F. D. Bona, and G. Rätsch, “Rna-seq read alignments with palmapper,” Current protocols in bioinformatics, pp. 11–6, 2010.

[25]. A. E. Jaffe, J. Shin, L. Collado-Torres, J. T. Leek, R. Tao, C. Li, Y. Gao, Y. Jia, B. J. Maher, T. M. Hyde, et al., “Developmental regulation of human cortex transcription and its clinical relevance at single base resolution,” Nature neuroscience, 2014.

[26]. T. Lappalainen, M. Sammeth, M. R. Friedländer, P. ACt Hoen, J. Monlong, M. A. Rivas, M. Gonzalez-Porta, N. Kurbatova, T. Griebel, P. G. Ferreira, et al., “Transcriptome and genome sequencing uncovers functional variation in humans,” Nature, 2013.

[27]. P. A. Combs and M. B. Eisen, “Low-cost, low-input rna-seq protocols perform nearly as well as high-input protocols,” PeerJ PrePrints, vol. 3.

[28]. F. Perez and B. E. Granger, “Ipython: a system for interactive scientific computing,” Computing in Science & Engineering, vol. 9, no. 3, pp. 21–29, 2007.

[29]. “Welcome to Apache Hadoop.” http://hadoop.apache.org, Aug. 2013.

[30]. J. Dean and S. Ghemawat, “Mapreduce: simplified data processing on large clusters,” Communications of the ACM, vol. 51, no. 1, pp. 107–113, 2008.

[31]. M. C. Schatz, B. Langmead, and S. L. Salzberg, “Cloud computing and the dna data race,” Nature biotechnology, vol. 28, no. 7, pp. 691–693, 2010.

[32]. L. D. Stein et al., “The case for cloud computing in genome informatics,” Genome Biol, vol. 11, no. 5, p. 207, 2010.

[33]. B. Langmead and S. L. Salzberg, “Fast gapped-read alignment with bowtie 2,” Nature methods, vol. 9, no. 4, pp. 357–359, 2012.

[34]. “GEUVADIS data access portal from the european nucleotide archive.” http://www.ebi.ac.uk/ena/data/view/ERP001942”uk/ena/data/view/ERP001942. Accessed: 2014-10-02.

[35]. R. Leinonen, R. Akhtar, E. Birney, L. Bower, A. Cerdeno-Tárraga, Y. Cheng, I. Cleland, N. Faruque, N. Goodgame, R. Gibson, et al., “The european nucleotide archive,” Nucleic acids research, p. gkq967, 2010.

[36]. T. Griebel, B. Zacher, P. Ribeca, E. Raineri, V. Lacroix, R. Guigó, and M. Sammeth, “Modelling and simulating generic rna-seq experiments with the flux simulator,” Nucleic acids research, vol. 40, no. 20, pp. 10073–10083, 2012.

[37]. “Rpkms for mrna sequencing data computed by GEUVADIS consortium.” http://www.ebi.ac.uk/arrayexpress/files/E-GEUV-3/GD660.TrQuantRPKM.txt.gz”ac.uk/arrayexpress/files/E-GEUV-3/GD660.TrQuantRPKM.txt.gz. Accessed: 2014-10-02.

[38]. Y. Liao, G. K. Smyth, and W. Shi, “The subread aligner: fast, accurate and scalable read mapping by seed-and-vote,” Nucleic acids research, vol. 41, no. 10, pp. e108–e108, 2013.

[39]. D. Kim, B. Langmead, and S. L. Salzberg, “Hisat: a fast spliced aligner with low memory requirements,” Nature methods, 2015.

[40]. F. Cunningham, M. R. Amode, D. Barrell, K. Beal, K. Billis, S. Brent, D. Carvalho-Silva, P. Clapham, G. Coates, S. Fitzgerald, et al., “Ensembl 2015,” Nucleic acids research, vol. 43, no. D1, pp. D662–D669, 2015.

[41]. W. J. Kent, A. S. Zweig, G. Barber, A. S. Hinrichs, and D. Karolchik, “Bigwig and bigbed: enabling browsing of large distributed datasets,” Bioinformatics, vol. 26, no. 17, pp. 2204–2207, 2010.

[42]. H. Li, B. Handsaker, A. Wysoker, T. Fennell, J. Ruan, N. Homer, G. Marth, G. Abecasis, R. Durbin, et al., “The sequence alignment/map format and samtools,” Bioinformatics, vol. 25, no. 16, pp. 2078–2079, 2009.

[43]. L. Collado-Torres, A. C. Frazee, M. I. Love, R. A. Irizarry, A. E. Jae, and J. T. Leek, “derfinder: Software for annotation-agnostic rna-seq differential expression analysis,” bioRxiv, p. 015370, 2015.

[44]. P. AC’t Hoen, M. R. Friedländer, J. Almlöf, M. Sammeth, I. Pulyakhina, S. Y. Anvar, J. F. Laros, H. P. Buermans, O. Karlberg, M. Brännvall, et al., “Reproducibility of high-throughput mrna and small rna sequencing across laboratories,” Nature biotechnology, 2013.

[45]. “Bowtie 2 manual.” http://bowtie-bio.sourceforge.net/bowtie2/manual.shtml. Accessed: 2015-08-08.

[46]. B. Langmead, C. Trapnell, M. Pop, S. L. Salzberg, et al., “Ultrafast and memory-efficient alignment of short dna sequences to the human genome,” Genome Biol, vol. 10, no. 3, p. R25, 2009.

[47]. W. J. Kent, C. W. Sugnet, T. S. Furey, K. M. Roskin, T. H. Pringle, A. M. Zahler, and D. Haussler, “The human genome browser at ucsc,” Genome research, vol. 12, no. 6, pp. 996– 1006, 2002.

[48]. “TopHat 2 manual.” http://ccb.jhu.edu/software/tophat/manual.shtml. Accessed: 2014-10-14.

[49]. “Amazon elastic compute cloud instance type information.” “http://aws.amazon.com/ec2/instance-types/”instance-types/. Accessed: 2014-10-16.

[50]. S. B. Montgomery, M. Sammeth, M. Gutierrez-Arcelus, R. P. Lach, C. Ingle, J. Nisbett, R. Guigo, and E. T. Dermitzakis, “Transcriptome genetics using second generation sequencing in a caucasian population,” Nature, vol. 464, no. 7289, pp. 773–777, 2010.

[51]. J. Harrow, A. Frankish, J. M. Gonzalez, E. Tapanari, M. Diekhans, F. Kokocinski, B. L. Aken, D. Barrell, A. Zadissa, S. Searle, et al., “Gencode: the reference human genome annotation for the encode project,” Genome research, vol. 22, no. 9, pp. 1760–1774, 2012.

[52]. “STAR manual.” http://github.com/alexdobin/STAR/blob/master/doc/STARmanual.pdf”pdf. Accessed: 2015-03-26.

[53]. P. G. Engström, T. Steijger, B. Sipos, G. R. Grant, A. Kahles, G. Rätsch, N. Goldman, T. J. Hubbard, J. Harrow, R. Guigó, et al., “Systematic evaluation of spliced alignment programs for rna-seq data,” Nature methods, 2013.

[54]. L. Collado-Torres, A. Nellore, A. Jae, J. Leek, and B. Langmead, “GEUVADIS expressed regions coverage matrix.” http://dx.doi.org/10.6084/m9.figshare.1405638.

[55]. “PyPy website.” http://pypy.org. Accessed: 2015-08-10.

[56]. “Amazon Elastic MapReduce (Amazon EMR).” http://aws.amazon.com/emr, Jan. 2014.

[57]. “Elephant Bird github repository.” http://github.com/kevinweil/elephant-bird/. Accessed: 2014-10-14.

[58]. M. Burset, I. Seledtsov, and V. Solovyev, “Analysis of canonical and non-canonical splice sites in mammalian genomes,” Nucleic Acids Research, vol. 28, no. 21, pp. 4364–4375, 2000.

[59]. N. Bansal, A. Blum, and S. Chawla, “Correlation clustering,” Machine Learning, vol. 56, no. 1-3, pp. 89–113, 2004.

[60]. N. Ailon, M. Charikar, and A. Newman, “Aggregating inconsistent information: ranking and clustering,” Journal of the ACM (JACM), vol. 55, no. 5, p. 23, 2008.

[61]. C. Bron and J. Kerbosch, “Algorithm 457: finding all cliques of an undirected graph,” Communications of the ACM, vol. 16, no. 9, pp. 575–577, 1973.

[62]. D. Eppstein, M. Löffer, and D. Strash, Listing all maximal cliques in sparse graphs in nearoptimal time. Springer, 2010.

[63]. “Bedgraph format specification.” http://genome.ucsc.edu/goldenpath/help/bedgraph.html” html. Accessed: 2014-10-26.

[64]. J. H. Bullard, E. Purdom, K. D. Hansen, and S. Dudoit, “Evaluation of statistical methods for normalization and differential expression in mrna-seq experiments,” BMC bioinformatics, vol. 11, no. 1, p. 94, 2010.

